# A Fasciclin 2 functional switch controls organ size in *Drosophila*

**DOI:** 10.1101/356196

**Authors:** Emma Velasquez, Jose A. Gomez-Sanchez, Emmanuelle Donier, Carmen Grijota-Martinez, Hugo Cabedo, Luis Garcia-Alonso

**Affiliations:** Instituto de Neurociencias CSIC-UMH, Universidad Miguel Hernandez. Sant Joan d’Alacant. Alicante 03550. Spain; Instituto de Investigación Sanitaria y Biomédica de Alicante (ISABIAL)-FISABIO. Sant Joan d’Alacant. Alicante 03550. Spain; Present address: Instituto de Investigaciones Biomedicas “Alberto Sols” CSIC-UAM, Universidad Autonoma de Madrid. Arturo Duperier 4, Madrid 28029. Spain

**Keywords:** Fasciclin 2, IgCAM, organ size, cell competition, imaginal disc, coupled MARCM, EGFR, Drosophila

## Abstract

How cell to cell interactions control local tissue growth to attain a species-specific pattern and organ size is a central question in developmental biology. The *Drosophila* Neural Cell Adhesion Molecule, Fasciclin 2 (*Drosophila* NCAM), is expressed during the development of neural and epithelial organs. Genetic mosaic analysis of Fasciclin 2 reveals two complementary and opposing functions during imaginal disc growth, a cell autonomous requirement to promote growth and an opposite non-cell autonomous function to restrain growth at high expression levels. This non-cell autonomous function is mediated by the Fasciclin 2 heterophilic-binding partners CG15630 and CG33543. We show that EGFR physically interacts with Fasciclin 2 and mediates both the cell autonomous and the non-cell autonomous function. We further show that EGFR activity in turn promotes the cell autonomous expression of Fasciclin 2. We suggest that the auto-stimulatory loop between EGFR and Fasciclin 2 operates until reaching a threshold where the Fasciclin 2 non-cell autonomous function counteracts the growth-promoting activity of the homophilic interaction to terminate imaginal disc growth. Accordingly, we have found that Fasciclin 2 limits imaginal disc growth by the end of larval development. Cellular integration of Fasciclin 2 autonomous and non-cell autonomous signaling from neighbor cells may be a key regulator component to orchestrate the rate of intercalary cell proliferation and the final size of an organ.

**Author Summary:** One of the key unsolved problems in Biology is how a species-specific size is attained during animal development. During development cells should compute the amount of intercalary tissue growth to stop cell proliferation when reaching a correct pattern and size. Classic studies demonstrated that local cell interactions are key in controlling organ growth to reach a correct size and pattern in vertebrates and invertebrates. We present evidence strongly suggesting that Fasciclin 2 (the ortholog of NCAM in *Drosophila*) functions as a growth level switch to control pattern and organ size. First, we use genetic mosaic analyses to show that Fasciclin 2 promotes organ growth in a cell autonomous manner. Then we show that Fasciclin 2 restrains growth at high expression levels in a non-cell autonomous manner, and that there is a requirement for Fasciclin 2 to limit growth by the end of larval development. This function is dependent on Fasciclin 2 heterophilic binding partners CG15630 and CG33543. The Epidermal Growth Factor receptor mediates both functional facets of Fasciclin 2 and its activity in turn increases Fasciclin 2 cell autonomous expression, suggesting the existence of a functional auto-stimulatory loop. We also show that the Epidermal Growth Factor receptor and Fasciclin 2 physically interact. Our results show that the amount of Fasciclin 2 between cells determines organ size by acting as an expression level switch for EGFR function, and suggest that other specific CAM interactions may integrate similar expression level switches acting as a code for cells to compute local growth in attaining a species-specific organ size and shape.

## Introduction

Morphogenesis involves the generation of organs with a species-specific size and shape. Cell proliferation is tightly controlled in rate and space during organ development to ensure a correct morphogenesis but, at the same time, its control is flexible enough to accommodate to perturbation. Control of growth occurs at the systemic, organ and tissue organization levels by the action of hormones, morphogens and cell interactions [1]-[4]. Despite the enormous progress achieved in identifying and characterizing the mechanisms involved in promoting or inhibiting growth, it remains a mystery how organs recognize when their local growth has reached the species-specific correct number of cells. Classic work demonstrated that local cell interactions are key in controlling intercalary cell proliferation to attain the final pattern and correct number of cells in vertebrate and invertebrate organs [5]. It is likely that cell-cell interaction mechanisms help cells compute the amount of local tissue growth, and operate upon growth signaling pathways to terminate cell proliferation when reaching a specific organ size [6]. Cell adhesion proteins of the immunoglobulin superfamily (IgCAMs) couple highly specific cell recognition and adhesion with the control of Receptor Tyrosine Kinase signaling [7], [8].

NCAM-type proteins function through homophilic adhesion and have been reported to act promoting the activation of the Fibroblastic Growth Factor Receptor (FGFR) and the Epidermal Growth Factor Receptor (EGFR) during axon growth [7], [8]. *Drosophila* conserves a homolog of the NCAM family of IgCAMs, Fasciclin 2 (Fas2), which is expressed in neural and non-neural tissues [9]-[11]. Fas2 was originally characterized as a homophilic IgCAM but it can also mediate heterophilic interactions with other IgCAMs [12]. Fas2 has been proposed to promote EGFR activity during axon growth [8]. In contrast, *fas2* loss-of-function (LOF) mutations have been found to cause derepression of the EGFR during retinal differentiation [13], and to produce *warts*-mediated cell over-proliferation in the follicular epithelium of the ovary [14].

*Drosophila* imaginal discs are epithelial organ precursors of the adult epidermis. Cell proliferation during imaginal disc growth is controlled by competitive cell interactions that help ensure the constancy of organ size and shape [15]. We have analyzed the role of Fas2 during imaginal disc growth using coupled-MARCM [16] and FLP-OUT genetic mosaics for LOF and gain-of-function (GOF) conditions. Our results show that Fas2 expression is required for growth in a dose-dependent and cell autonomous manner. Clones of cells devoid of Fas2 grow less than normal and express increased levels of JNK activity. Reducing the competitive character of the normal cells outside of the clone reverts JNK expression in the Fas2-deficient clones, indicating that activation of the JNK pathway is a secondary effect of the competition between Fas2-deficient slow cells and their normal neighbors. Neither inhibition of JNK signaling nor suppression of apoptosis in the Fas2-deficient cells rescues the size of the clones, demonstrating that their small size is due to reduced cell proliferation. Analysis of the mitotic index in Fas2-deficent clones confirms this inference.

Fas2 promotes EGFR activity during imaginal disc growth, as revealed by the reduced ppERK expression in Fas2-deficient cells and the genetic interactions between *fas2* LOF conditions and EGFR mutations. Fas2-deficient clone rescue analysis shows that EGFR and its effectors Ras and Raf specifically act to mediate Fas2 function, while the Hippo signaling pathway is indirectly involved downstream of JNK activation. Furthermore, co-immunoprecipitation experiments show that Fas2 physically binds EGFR in cultured cells. Interestingly, EGFR function in turn promotes cell autonomous expression of Fas2, suggesting the existence of a self-stimulatory feedback loop between Fas2 expression and EGFR activity.

GOF analysis of Fas2 during imaginal disc growth shows it can restrain cell proliferation in a non-cell autonomous manner, causing a dose-dependent reduction of organ size at high expression levels due to a reduction of EGFR function. We show that this functional facet of Fas2 is mediated by its heterophilic binding partners CG15630 and CG33543. These IgCAMs are expressed in all epithelial cells, although their domain of expression does not exactly match that of Fas2 in the cell membrane. CG15630 and CG33543 are both required for growth and their over-expression causes a growth deficit, thus paralleling the functionality of Fas2. Taken together, our results suggest a scenario where low and moderate levels of Fas2 promote EGFR activity in each epithelial cell during the growth of imaginal discs, which in turn enhances Fas2 expression. This positive cell autonomous feedback loop between Fas2 and the EGFR may progress until being counteracted by the interaction mediated by CG33543 and CG15630 at high Fas2 expression levels. Accordingly, we show that partial mild inhibition of Fas2 expression during the last day of larval development causes some extra-growth, demonstrating that both Fas2 functional facets act in concert during imaginal disc development. Thus, the amount of Fas2 between cells determines whether to grow or not to grow, acting as an expression level growth switch in development.

## Results

### Loss of Fas2 causes a cell autonomous deficit of growth in imaginal discs

All epithelial cells in imaginal discs express Fas2 in a dynamic pattern [10] (Fig. 1A, B). The maximum expression corresponds to differentiating or pre-differentiating structures, like veins, proneural clusters, sensory organ precursor cells, and the Morphogenetic Furrow and photoreceptors in the eye imaginal disc. Despite these peaks of highest expression, undifferentiated proliferating epithelial cells also express Fas2 at lower levels (Fig. 1 A, B). Individuals lacking Fas2 are lethal but whole *fas2*^*–*^ null organs can develop and differentiate epidermis in gynandromorphs [10] (Fig. S1A). We analyzed *fas2* hypomorphic LOF conditions generated by either hypomorphic allele combination or by restricted *RNAi* expression in the wing or the eye imaginal disc. Both, the hypomorphic mutant combination and the expression of *fas2*^*RNAi*^ in the wing imaginal disc driven by *MS1096-GAL4* caused size reductions of the whole adult wing (Fig. 1C, G). Expression of *fas2*^*RNAi*^ in the posterior compartment of the wing imaginal disc using the *engrailed-GAL4* driver caused a specific reduction in the size of the compartment in late 3^rd^ instar larva (Fig. 1E). Expression of *fas2*^*RNAi*^ in *eyeless*-driven FLP-OUTs caused lethality in pupa after head eversion. Some escaper individuals managed to eclose as active adult flies showing a reduced head size (Fig. 1D). Organ size reduction was evident in the eye-antenna imaginal discs of these larvae before puparium formation (Fig. 1D). Organ size reduction was due to a lower number of cells, and not to a reduced cell size, as revealed by either the normal spacing of trichomes in the adult wing (which mark each single epidermal cell) (Fig. 1C) or the cell profiles in imaginal discs and pupal wings (Fig. 1E, F).

**Fig. 1.**
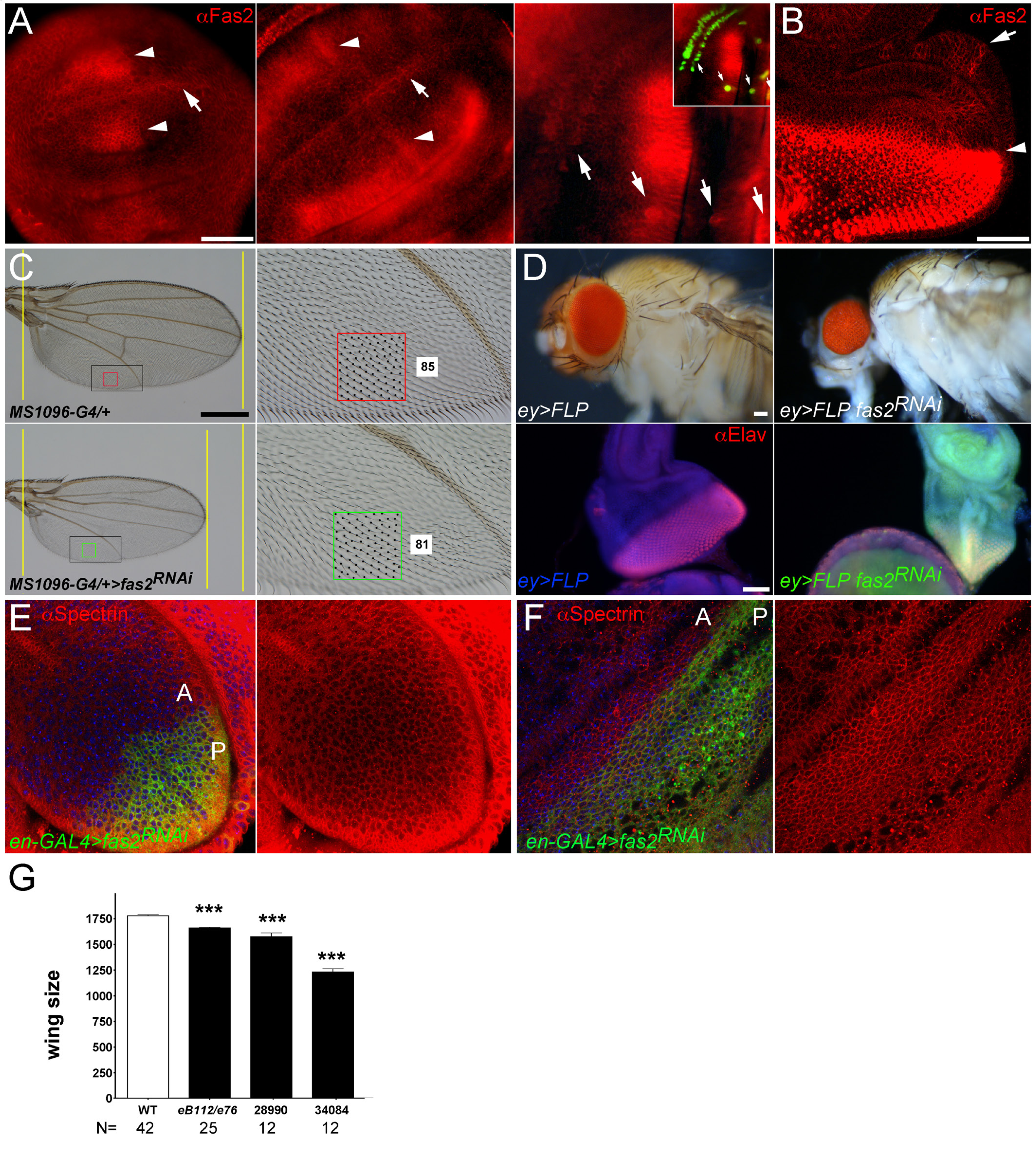
Expression and requirement of Fas2 in imaginal discs. (A) Fas2 expression (MAb-1D4 antibody) in early, middle and late 3^rd^ instar larva wing discs. Note the changing levels of Fas2 expression at both sides (arrowheads) of the D/V compartment border (arrow) in early and middle 3^rd^ instar wing discs (left and center). In late 3^rd^ instar wing imaginal discs (at right) sensory organ precursor cells (arrows, marked with *neuralized-LacZ* in inset) express high levels of the protein. Bar: 50 μm. (B) Late 3^rd^ instar larva eye-antenna imaginal disc. All cells express Fas2 in a dynamic pattern. The highest levels are found at the Morphogenetic Furrow (arrowhead), in the differentiating retina posterior to it and in the Ocellar prospective region (arrow). Bar: 50 μm. (C) Expression of *UAS-fas2*^*RNAi*^ (#34084) in the wing disc under the control of the *MS1096-GAL4* insertion (doubly heterozygous females) causes a reduction of wing size (compare the two left panels), and the loss of cross-veins. The size reduction is due to a lower number of epidermal cells in the *fas2*^*RNAi*^ wing (compare the two right panels for the number of trichomes, each one representing a single wing cell). The spacing and polarity of trichomes is normal in the *fas2*^*RNAi*^ wing (85 vs. 81 cells inside the color squares). Bar: 500 μm. (D) Expression of *UAS-fas2*^*RNAi*^ (#34084) in the eye-antenna imaginal disc in *eyeless*-driven FLP-OUTs (*ey>FLP*) causes a reduction of the size of the whole head (doubly heterozygous females, upper panels). The reduction of size is also evident in 3^rd^ instar larva eye-antenna imaginal discs (lower panels). At left, a control *ey-FLP; CyO/+; UAS-fas2*^*RNAi34084*^*/+* sibling eye-antenna imaginal disc stained with anti-Elav to reveal the neurons in the differentiating retina. At right, an *ey-FLP; ActinFRTy*^*+*^*FRT-GAL4 UAS-GFP/+; UAS-fas2*^*RNAi34084*^*/+* eye-antenna imaginal disc. Note the severe reduction in size caused by the expression of *fas2*^*RNAi*^. Bar: 50 μm. (E) Expression of *UAS-fas2*^*RNAi*^ (#34084) under the control of the *engrailed-GAL4* (*en-GAL4*) driver (*en-Gal4>fas2*^*RNAi*^) causes a reduction in the size of the posterior compartment (P) of the wing imaginal disc. The shape and size of the cells is not affected by the expression of the *fas2*^*RNAi*^ (compare cells in the anterior, A, and posterior, P, compartments). (F) During pupal development, expression of *UAS-fas2*^*RNAi*^ (#34084) in the P compartment of the wing under the control of the *en-GAL4* driver does not alter cell size or shape. (G) Quantitative analysis of wing area in *fas2* LOF mutant combinations. WT controls are *MS1096-GAL4* heterozygous females. The hypomorphic *fas2*^*eB112*^*/fas2*^*e76*^ combination causes a moderate but extremely significant reduction of wing size, while the expression of *fas2*^*RNAi*^, RNAi #28990 or RNAi #34084, in *MS1096-GAL4* heterozygous females cause a more dramatic size reduction. Wing size is wing area in μm^2^/10^3^.

To study the cellular requirement of Fas2 in the null condition, we generated *fas2*^*–*^ null cell clones during the 1^st^ and 2^nd^ larval stages of larval development using the coupled-MARCM technique. Fas2-deficient clones were extremely reduced in size or absent compared to wild type clones, or their own control twins in 3^rd^ instar larva wing and leg imaginal discs (Fig. 2A, E; Fig. S1B, D), eye disc (Fig. 2C), pupal wing (Fig. 2D), and the adult structures derived from imaginal discs (Fig. S1C, E, F), but were more normal in the larval brain (Fig. 2C) and the adult abdomen (Fig. S1C). In some cases, *fas2*^*+*^ twin control clones in imaginal discs or their derivatives were exceptionally large (Fig. S1B, E). The size of pupal wings or imaginal discs bearing coupled-MARCM *fas2*^*–*^ clones was the same than WT coupled-MARCM controls (Fig. 2D, F), indicating that the loss of growth produced by the *fas2*^*–*^ clones was compensated by the growth of the normal cells to attain a correct final organ size. Analysis of the mitotic index in coupled-MARCM *fas2*^*–*^ clones revealed a significant decrease compared to WT or their control twin clones (Fig. 2G; Fig. S1D). Interestingly, the normal neighbor cells closest to *fas2*^*–*^ clones displayed a reduced mitotic index as well, strongly suggesting that it is the loss of Fas2 homophilic binding what hinders cell proliferation. The growth deficit of *fas2*^*–*^ clones was rescued by the expression of either the GPI-anchored (Fas2^GPI^), the trans-membrane (Fas2^TM^) isoform of the protein, or both at the same time (Fig. 2B, C). Thus, *fas2*^*–*^ cells can proliferate and differentiate normal epidermis in whole *fas2*^*–*^ organs [10], but they are hampered to do so when developing along with other cells expressing Fas2 in the same imaginal disc. This behavior of *fas2*^*–*^ cells in clones is symptomatic of cell competition. It strongly suggests that slow proliferating Fas2-deficient cells are out-competed by normal neighbor cells expressing Fas2. To confirm the presence of cell competition, we induced Fas2-deficient clones using the Minute method [17], which slows the proliferation rate of all the cells in the organ with the exception of those in the clone. *fas2*^*–*^ *Minute*^*+*^ clones were significantly larger than *fas2*^*–*^ clones (Fig. S1G-I), thus confirming that Fas2 is required for normal cell proliferation and competition in imaginal discs. *fas2*^*–*^ *Minute*^*+*^ clones displayed a strong tendency to intermingle in the *Minute*^*–*^*/+* heterozygous genetic background (Fig. S1G).

**Fig. 2.**
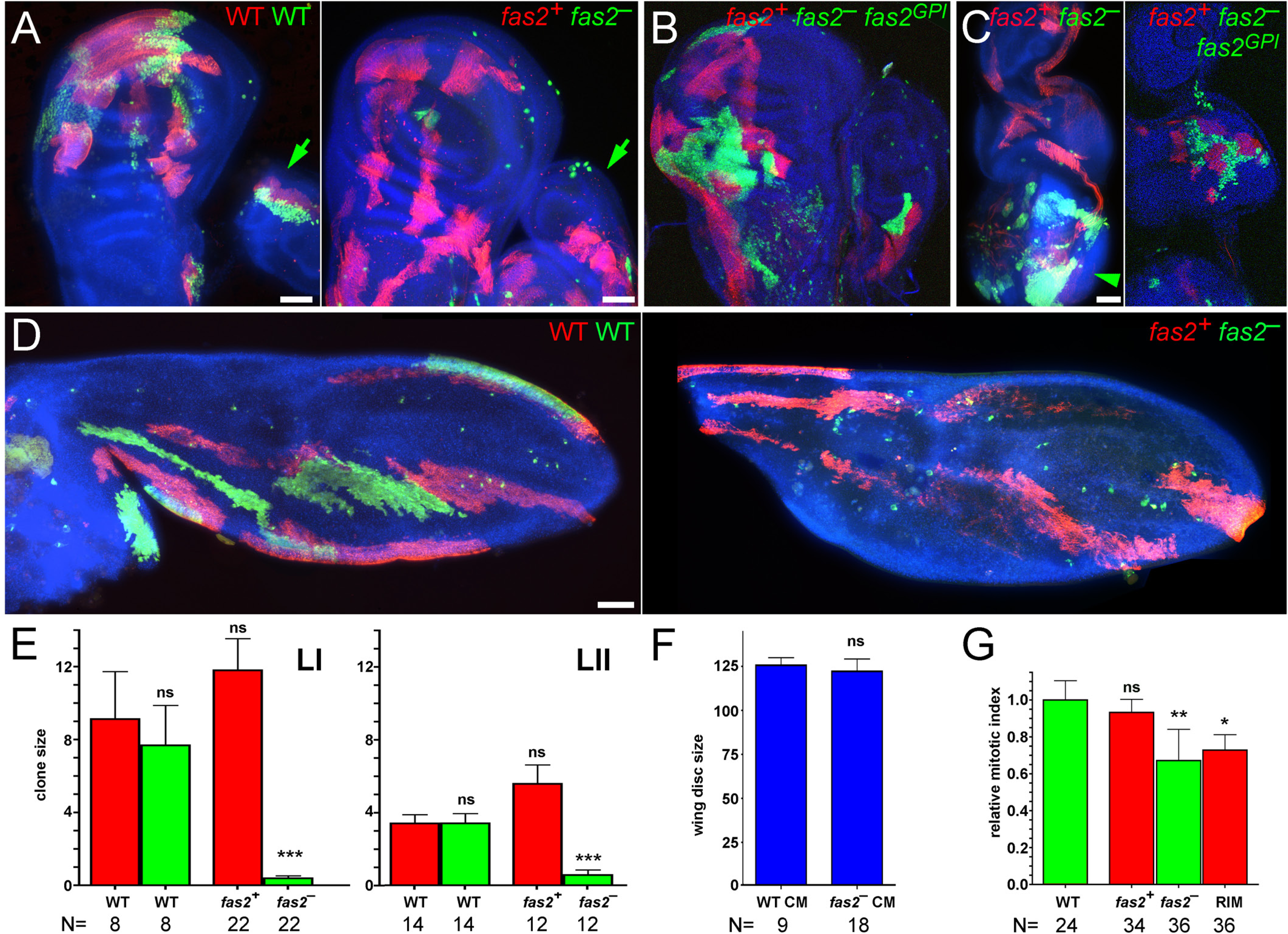
Cell autonomous requirement of Fas2 for cell proliferation in imaginal discs. (A) Left, coupled-MARCM analysis in WT. Wing disc twin clones labeled with GFP (WT) and mtdTomato (WT) induced in 1^st^ instar larva. Bar: 50 mm. Right, coupled-MARCM analysis of the *fas2* null condition (*fas2*^*eB112*^). Wing disc *fas2*^*–*^ null clones (labeled with GFP) induced in 1^st^ instar larva are missing or extremely small compared with their control twins (labeled with mtdTomato). Leg disc *fas2*^*–*^ clones are also missing or reduced in size compared to WT (arrows). (B) Coupled-MARCM analysis of *fas2* null clones rescued by the expression of *UAS-fas2*^*GPI*^isoform. (C) In the eye-antenna imaginal disc, *fas2*^*–*^ clones display the exact same behavior than in the wing disc, and are also rescued by the expression of the GPI-linked isoform of Fas2. Note that the *fas2*^*–*^ clones do have a more normal size in the brain lobe (arrowhead). Bar: 50 μm. (D) Left, coupled-MARCM WT twin clones in pupal wings induced in 1^st^ instar larva. Bar: 50 mm. Right, coupled-MARCM analysis of the *fas2*^*–*^ null condition in pupal wings. Fas2-deficient clones (*fas2*^*–*^ GFP) induced in 1^st^ instar larva are either missing or extremely small compared with their control twins (*fas2*^*+*^ mtdTomato). (E) Quantitative analysis of coupled-MARCM clones in the wing disc. Wing imaginal disc *fas2*^*–*^ null clones (*fas2*^*–*^ GFP, green bars) induced in 1^st^ (LI) and 2^nd^ instar larva (LII) are extremely small compared with either their internal control twins (*fas2*^*+*^ mtdTomato, red bars) or with coupled-MARCM WT twin clones (WT GFP, green bars, and WT mtdTomato, red bars). Note that the internal control *fas2*^*+*^ twin clones, which are growing in the heterozygous *fas2*^*–*^/+ genetic background, display a clear tendency to be larger than the WT coupled-MARCM controls. Clone mean size in μm^2^/10^3^. (F) Wing discs harboring coupled-MARCM *fas2*^*–*^ null clones (*fas2*^*–*^ CM) induced in 1^st^ and 2^nd^ instar larva have the same size than control coupled-MARCM (WT CM) discs. (G) Comparison of mitotic index in *fas2*^*eB112*^ null clones and their coupled-MARCM control twins. Twin clones display a similar mitotic index to WT clones. *fas2*^*eB112*^ null clones and the rim of normal cells around the clone show a significant reduction in the mitotic index.

**Fig. S1. Fas2 requirement in imaginal discs**

### Fas2 insufficiency indirectly causes a cell competition-dependent activation of the JNK pathway

Apoptosis of slow proliferating cell clones is a consequence and integral part in the process of cell competition. However, *fas2*^*–*^ cell clones were able, or at least some of them, to evade apoptosis and differentiate in adult epidermis (Fig. S1F, J). To study the contribution of apoptosis to the fas2 phenotype, we combined the *fas2*^*–*^ genotype with different genetic conditions that suppress cell death. Expression of the Drosophila-Inhibitor-of-Apoptosis, DIAP1 (*UAS-diap*) [18] in the *fas2*^*–*^ clones produced a slight but statistically significant normalization of clone size in the adult (Fig. S1K). *Df(3L)H99/+*, which has a very strong anti-apoptotic effect during cell competition [19], also caused a slight but statistically significant increase in the size of the *fas2*^*–*^ clones (Fig. S1K). On the other hand, we did not detect any significant modification of the phenotype expressing P35 in the clones (Fig. S1K).

The JNK signaling pathway controls apoptosis and has been proposed to interact with Fas2 during neural differentiation in pupa. [20]. To test the involvement of this signaling pathway in the *fas2* phenotype during imaginal disc growth, we induced the inhibition of JNK (Bsk) activity in the wing of *MS1096-Gal4/+; UAS-fas2*^*RNAi*^*/UAS-bsk*^*RNAi*^ and in the eye disc of *eyeless*-driven *UAS-fas2*^*RNAi*^*/UAS-bsk*^*RNAi*^ FLP-OUT individuals. Wings and eyes respectively only displayed a partially corrected size, and they continued to be significantly smaller than their *MS1096-Gal4/+; UAS-bsk*^*RNAi*^*/+* or *eyeless-* driven *UAS-bsk*^*RNAi*^ FLP-OUT controls (Fig. 3A, B; Fig S1L). On the other hand, coupled-MARCM *fas2*^*–*^ *bsk*^*RNAi*^ clones continued being much smaller than their control twins (Fig. 3C). Analysis of the activity of JNK (phosphorylated-JNK) in *en-GAL4 UAS-fas2*^*RNAi*^ revealed only scattered cells in the posterior and anterior compartments (Fig. 3D), and expression of cleaved Caspase 3 did not show obvious signs of apoptosis (Fig. 3E).

**Fig. 3.**
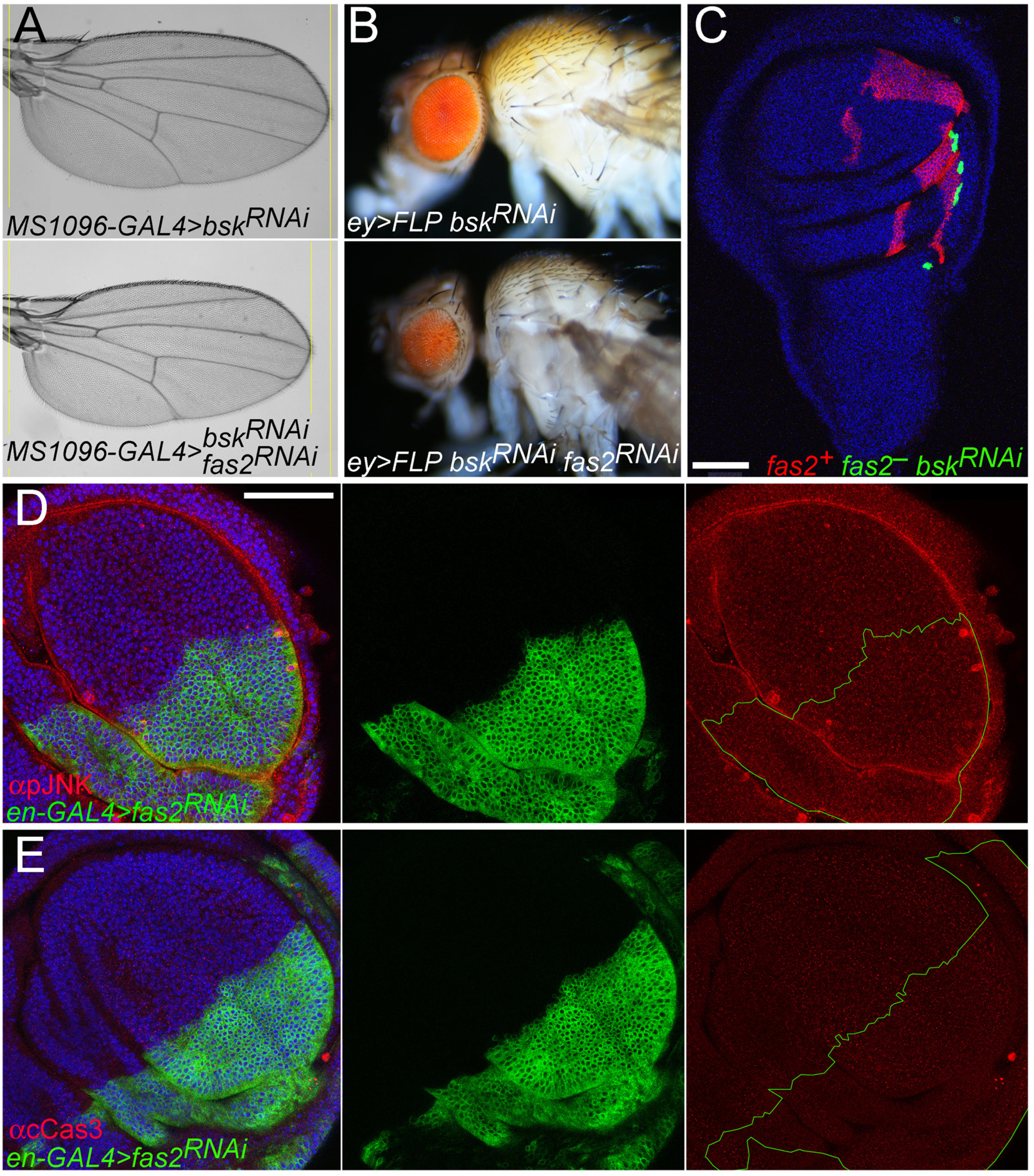
Analysis of JNK signaling in fas2 LOF conditions. (A) Inhibition of the JNK signaling pathway using *UAS-bsk*^*RNAi*^ (#32977) ameliorates but does not rescue the reduced wing size caused by the expression of *UAS-fas2*^*RNAi*^ (#34084) under the control of the *MS1096-GAL4* driver (double heterozygous females). (B) Expression of *UAS-bsk*^*RNAi*^ (#57035) ameliorates but does not rescue the reduced eye phenotype produced by expression of *UAS-fas2*^*RNA*i^ (#34084) in *ey*-driven FLP-OUT (*ey>FLP*) individuals. (C) Coupled-MARCM *fas2*^*eB112*^ null clones expressing *UAS-bsk*^*RNAi*^ (#32977) (GFP) display the typical reduced size compared to their normal twin controls (mtdTomato). Bar: 50 μm. (D) The posterior compartment of an *en-GAL4/+* wing disc expressing *UAS-fas2*^*RNAi*^ (#34084) shows scattered cells expressing phosphorylated-JNK. Bar: 50 μm. (E) *en-GAL4/+; UAS-fas2*^*RNAi#34084*^*/+* wing discs do not show obvious signs of apoptosis using an anti-cleaved Caspase3 antibody.

To further analyze the activity of the JNK pathway in the Fas2-deficient cells, we monitorized the expression of the JNK pathway reporter *puckered-LacZ* (*puc-LacZ*) [21]. Puc is a negative feedback repressor of the JNK pathway, and the *puc-LacZ* insertion causes a null mutation in the *puc* gene. Expression of *UAS*-*fas2 RNAi* in the posterior compartment of the wing using the *en-GAL4* driver caused a reduction of compartment size due to a reduced number of cells, but no widespread signs of apoptosis (Fig. 3D, E). The introduction of the heterozygous *puc-LacZ* insertion in this genotype revealed a strong expression of the reporter (Fig. 4A), indicating that *fas2*^*RNAi*^-expressing cells had an enhanced activity of the JNK signaling pathway in combination with the 50% reduction of the *puc* gene dosage. This reduction in half of the normal dose of Puc caused widespread apoptosis, increased expression of phosphorylated-JNK (including some Fas2-normal neighbor cells) and alterations in the Anterior-Posterior compartment border (Fig. 4B, C). Strong expression of *puc-LacZ* and concomitant increased apoptosis was also found in the *fas2*^*–*^ cells of MARCM Fas2-deficient clones (Fig. 4D). *puc-LacZ* expression, apoptosis and clone growth was reverted to normal by the simultaneous expression of the Fas2^GPI^ isoform in the clones (Fig. 4E), showing the extracellular part of Fas2 is sufficient to revert JNK activation and support the function of Fas2 on cell proliferation in imaginal discs. To determine if the JNK increased activity in the Fas2-deficient clones reflected a direct or an indirect function of Fas2 on controlling this signaling pathway, we analyzed *puc-LacZ* expression in *fas2*^*—*^ *Minute*^*+*^ clones. Clone growth was normal and LacZ expression only sporadic in *fas2*^*–*^ *M*^*+*^ *puc-LacZ/+* control clones rescued by constitutive expression of a *hs-fas2*^*TM*^ insertion (Fig. 4F), while expression of LacZ was completely abolished and growth reduced in the *fas2*^*–*^ *Minute*^*+*^ *puc-LacZ/+* clones (Fig. 4G). This result shows that JNK pathway activity in the *fas2*^*–*^ cells indirectly results from the cell competition process with the fast proliferating cellular background. In addition, these *fas2*^*–*^ *M*^*+*^ *puc-LacZ/+* clones were not able to grow as much as in a Puc-normal background (Fig. 4G; Fig. S1G). Since Puc is a specific feedback repressor of the JNK pathway [22] and the *puc-LacZ* insertion causes a mutation in the *puc* gene, our data together suggest that Puc expression is critical to balance JNK derepression to allow for Fas2-deficient cell survival and growth (see below). In summary, Fas2-deficient cells grow less than normal and can be out-competed by Fas2-normal neighbor cells, although some can survive to differentiate as adult epidermis by the compensatory action of the JNK-Puc regulatory loop.

**Fig. 4.**
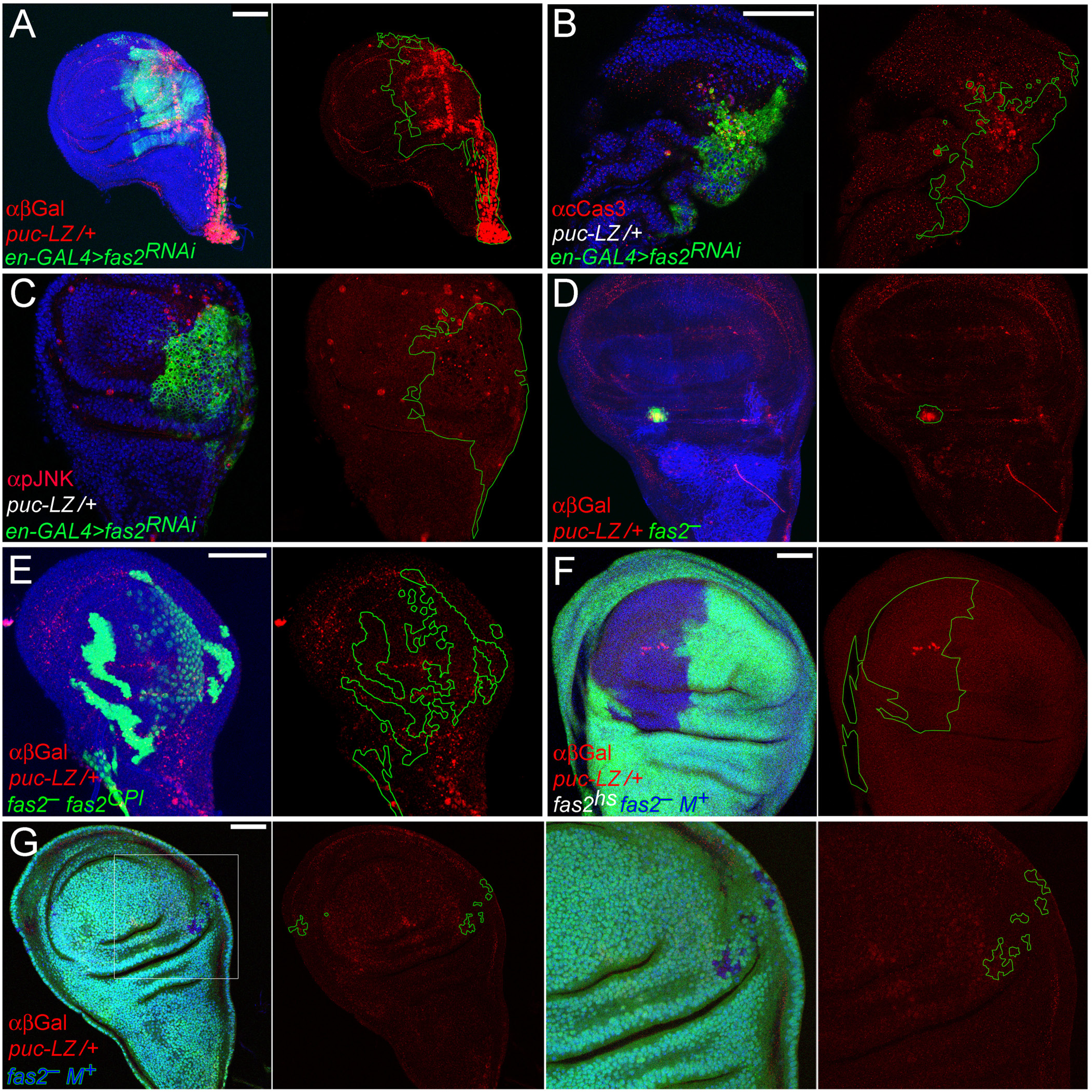
Activation of Puc-dependent JNK signaling in Fas2-deficient cells is indirect through cell competition. (A) Posterior compartments deficient for Fas2 in *en-GAL4/+; UAS-fas2*^*RNAi#34084*^*/puc-LacZ* individuals display strong expression of the reporter, indicating a strong enhancement of JNK activity. Bar: 50 μm. (B) Posterior compartments deficient for Fas2 display widespread apoptosis (staining with anti-cleaved Caspase 3 antibody) and the breakup of the compartment border. (C) The same individuals show strong derepression of activated-JNK (staining with an anti-phospho-JNK antibody). Note that phosphorylated-JNK occurs non-cell autonomously in the anterior compartment as well as in the posterior compartment. (D) MARCM *fas2*^*eB112*^; *puc-LacZ/+* cell clones display strong expression of the reporter, indicating derepression of the JNK signaling pathway. (E) Expression of the Fas2^GPI^ isoform in MARCM *fas2*^*eB112*^; *puc-LacZ/UAS-fas2*^*GPI*^ cell clones rescue the activation of JNK, indicating that the extracellular domain of Fas2 is sufficient to sustain Fas2 function. (F) The Minute technique allows *fas2*^*eB112*^; *hs-fas2*^*TM*^*/puc-LacZ* (*M*^*+*^) control cell clones to proliferate more than neighbor *Minute*^*–*^*/+* cells (labeled with *Ubi-GFP*). Only sporadic cells show activation of the JNK pathway. (G) Reducing the proliferation rate of the cellular background (labeled with *Ubi-GFP*) with the Minute technique rescues JNK derepression in *fas2*^*eB112*^; *puc-LacZ/+* cell clones. Note that the simultaneous deficit of Fas2 and half the dosage of Puc prevents the cell clones to overgrow (compare with the same clones rescued with *hs-fas2*^*TM*^ in F)

### EGFR mediates the cell autonomous function of Fas2 in epithelial cell proliferation

During axon growth Fas2 promotes EGFR function [8]. In contrast, during retinal differentiation *fas2* LOF conditions cause a derepression of EGFR function [13]. We analyzed the level of activated-MAPK expression (di-phosphoERK, a reporter of EGFR activity) in *eyeless*-driven (*ey>FLP*) *fas2*^*RNAi*^ FLP-OUT imaginal discs. These Fas2-deficient eye imaginal discs displayed a reduced level of activated-MAPK expression compared to their control siblings, or tissue of the same individual without *ey* expression (Fig. 5A), consistent with a reduced EGFR activity. Moreover, *fas2* LOF conditions displayed an adult phenotype reminiscent of *Egfr* and *Ras* LOF conditions (including reduced size, loss of ocelli and bristles, and loss of cross-veins in the wing) (Fig. 1C, D; Fig. S2A-D). The *fas2* LOF phenotype is reciprocal to that of *Egfr* GOF conditions (Fig. S2D). In addition, *fas2* acute GOF conditions (but not chronic conditions, see below) caused the differentiation of extra-veins, which resembled the typical phenotype of *Egfr* GOF conditions in the wing (Fig. S2C). These results suggest that Fas2 promotes EGFR activity during imaginal disc growth. Indeed, a 50% reduction of the *Egfr* dosage (which does not cause any phenotypic alteration on its own) enhanced the phenotype of the *fas2*^*eB112*^*/fas2*^*e76*^ hypomorphic mutant combination (Fig. 5 E), showing that Fas2 function is dependent on the dosage level of EGFR function. On the other hand, double mutant combinations between *Egfr* and *fas2* strong LOF conditions displayed an epistatic behavior (Fig. 5E). Thus, the genetic combination between *Egfr* hypomorphic alleles and the stronger LOF condition for *fas2* (*Egfr*^*top1*^*/Egfr*^*top1*^; *fas2*^*RNAi*^*/+* and *Egfr*^*top1*^*/Egfr*^*top2W74*^; *fas2*^*RNAi*^*/+*) displayed a wing size close to the *fas2* combination alone (*fas2*^*RNAi*^*/+*), and very different to the expected value for an additive phenotypic interaction (Fig. 5E). The results are strongly consistent with a positive interaction of Fas2 and EGFR in the same signaling pathway during imaginal disc growth. Therefore, Fas2 function during imaginal disc growth seems just opposite to its reported function during retinal differentiation [13].

**Fig. 5.**
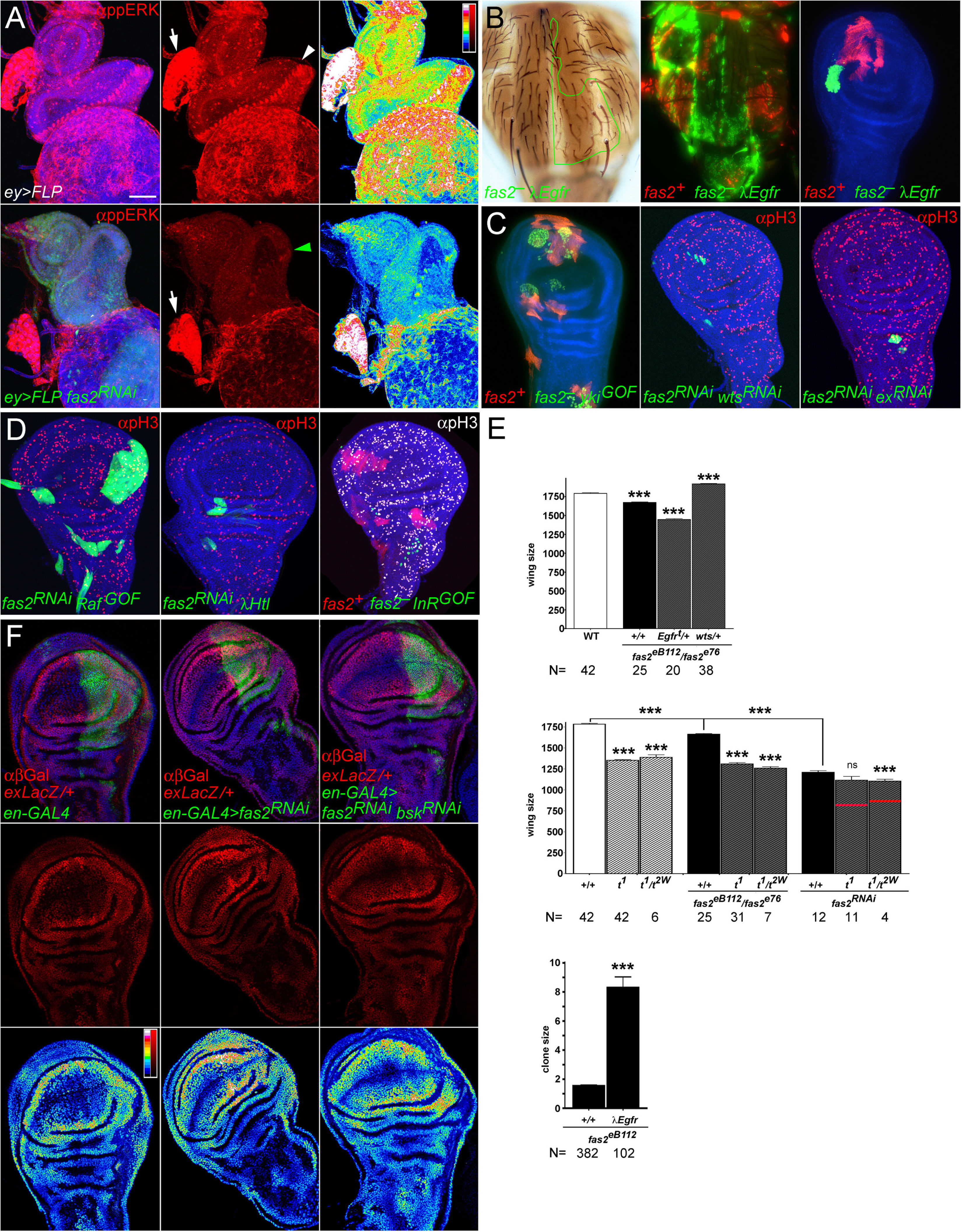
The EGFR signaling pathway mediates the Fas2 cell autonomous function to promote imaginal disc growth. (A) Eye imaginal discs deficient for Fas2 display reduced levels of activated-MAPK. Strong activated-MAPK expression is evident in the Morphogenetic Furrow of the eye disc (arrowhead) and the Ring Gland (arrow) of controls (CyO balancer siblings, top panels). *ey*-driven FLP-OUT eye discs expressing *UAS-fas2*^*RNAi*^ *(#34084)* (green arrowhead, bottom panels) show a strong reduction of activated-MAPK (anti-ppERK antibody). Note that the eye disc (where *ey-FLP* is expressed, labeled with UAS-GFP) displays a strong reduction in activated-MAPK expression compared with the Ring Gland of the same individual (which does not express ey-FLP, no GFP labeling, arrow). The right panels show a LUT color coded representation of activated-MAPK signal levels. Bar: 50 μm. (B) Rescue of the *fas2*^*–*^ phenotype by activated-EGFR *(λEGFR*) in adult (left), late pupa (middle) and the wing imaginal disc. At left, an adult *fas2*^*eB112*^ *λEgfr* MARCM clone in the notum marked with *yellow* and *forked* (outlined in green) showing a rescued size and extra bristles. Middle, *fas2*^*eB112*^ *λEgfr* coupled-MARCM clones in the notum (GFP) displaying a rescued size similar to their normal *fas2*^*+*^ control twins (labeled with mtdTomato). Right, *fas2*^*eB112*^ *λEgfr* coupled-MARCM clone and control twin (labeled with mtdTomato) in the wing disc. (C) Rescue of the *fas2* clone phenotype by Yki over-expression, but not by Ex or Wts inhibition. Right, coupled-MARCM *fas2*^*eB112*^ clones over-expressing Yki (*fas2*^*eB112*^; *UAS-yki/+*, labeled with GFP) and their *fas2*^*+*^ control twins (labeled with mtdTomato) in the wing disc. Middle, 2^nd^ instar larva *UAS-fas2*^*RNAi#34084*^ *UAS-wts*^*RNAi#37023*^ FLP-OUT clones (labeled with GFP). Right, 1^st^ instar larva *UAS-fas2*^*RNAi#34084*^ *UAS-ex*^*RNAi#34968*^ FLP-OUT clone (labeled with GFP). (D) Expression of activated-Raf (*UAS-Raf*^*GOF*^), but not of activated-FGFR (*UAS-λHtl*) or activated-InR (*UAS-InR*^*GOF*^), rescues *fas2* clone growth. Left, 2^nd^ instar larva FLP-OUT *UAS-fas2*^*RNAi#34084*^ *UAS-Raf*^*GOF*^*/+* clones. Middle, 2^nd^ instar larva FLP-OUT *UAS-fas2*^*RNAi#34084*^ *UAS-λHtl/+* clone. Right, 2^nd^ instar larva coupled-MARCM of *fas2*^*eB112*^; *UAS-InR*^*DEL*^*/+* clones (GFP, absent) and their control twins (mtdTomato). A similar result was obtained with the expression of *UAS-InR*^*R418P*^. (E) Top, the *fas2* wing phenotype is sensitive to the dosage of *EGFR* and *wts* genes. *fas2*^*eB112*^*/fas2*^*e76*^ wings show a reduction in size which is enhanced by a reduction of 50% in the dosage of *EGFR* (*Egfr*^*top2w74*^*/+*), and suppressed by a reduction of 50% in the dosage of *wts* (*wts*^*X1*^*/+*) (WT and *fas2*^*eB112*^*/fas2*^*e76*^ as in Fig. 1G). Middle, Functional interactions between *fas2* and *Egfr* LOF mutations. The size of *fas2*^*eB112*^*/fas2*^*e76*^ and *fas2*^*RNAi34084*^ (*MS1096-GAL4* driven) adult wings (females, black bars) is significantly smaller than WT (white bar, same as in Fig. 1G). Two *Egfr* hypomorphic combinations (*Egfr*^*top1*^*/Egfr*^*top1*^, *t*^*1*^, and *Egfr*^*top1*^*/Egfr*^*top2W74*^, *t*^*1*^*/t*^*2W*^; striped white bars) display a wing size smaller than *fas2*^*eB112*^*/fas2*^*e76*^ and similar to that caused by the expression of *UAS-fas2*^*RNAi34084*^. The wing size of the double mutant combinations of *fas2*^*eB112*^*/fas2*^*e76*^ with the *Egfr* allelic combinations is similar to the *Egfr* mutants (striped grey bars). The stronger *UAS-fas2*^*RNAi34084*^ condition shows an epistatic interaction with the *Egfr* combinations. The combination with *Egfr*^*top1*^*/Egfr*^*top2W74*^ shows a significative enhancement, but far from the value expected for an additive effect of the fas2 and Egfr phenotypes (red line). All individuals are female. Bottom, *y fas2*^*eB112*^ *f*^*36a*^ MARCM 2^nd^ instar larva clone size in adults is rescued by activated EGFR (*UAS-λEgfr/+*) (size is number of *y f*^*36a*^ bristles). Wing size is wing area in μm^2^/10^3^. (F) Reduction of Fas2 function by the expression of *UAS-fas2*^*RNAi*^ *(#34084)* under the control of the *en-GAL4* driver causes the over-expression of the *ex-LacZ* reporter. Left panels show a control *en-GAL4/ex-LacZ* wing disc. Middle panels show an *en-GAL4/ex-LacZ; UAS-fas2*^*RNAi34084*^*/+* wing disc (note the higher expression in the P compartment extending into the A compartment). Right panels show an *en-GAL4/ex-LacZ; UAS-fas2*^*RNAi34084*^*/UAS-bsk*^*RNAi#32977*^. Note the reversion in the expression of the *ex-LacZ* reporter. Bottom panels show a color coded LUT representation of signal intensity for LacZ.

**Fig. S2. *fas2* LOF and acute GOF phenotypes are reminiscent of *Egfr* LOF and GOF mutations respectively**

To further analyze the specificity of the interaction between Fas2 and the EGFR, and to identify other effectors that may mediate the Fas2 function in imaginal discs, we studied the capacity of different GOF or LOF conditions for proteins involved in growth signaling to suppress or enhance the phenotype of *fas2*^*–*^ clones (Fig. S2E). Activated-EGFR (*UAS-λEgfr*) rescued the size of *fas2*^*–*^ clones in imaginal discs, pupa and adult (Fig. 5B). Effectors of the EGFR signaling pathway: activated-Ras (*UAS-Ras*^*V12*^), activated-Raf (*UAS-Raf*^*GOF*^), active-PI3K (*UAS-Dp110*) (Fig. 5D; Fig. S2E) and over-expression of Yki (*UAS-Yki*) (Fig. 5C; Fig. S2E) also produced a significant normalization of *fas2*^*–*^ MARCM clones in the adult and imaginal discs. Therefore, the results indicate that the EGFR and its signaling pathway effectors function downstream of Fas2 during imaginal disc growth. In contrast, activated-FGFR (*UAS-λhtl*), activated-Insulin receptor (*UAS-InR*^*DEL*^, *UAS-InR*^*418P*^), over-expression of Src (*UAS-Src42A*) and activated-Notch (*UAS-N*^*INTRA*^) were unable to produce a significant effect in adult or imaginal disc clones (Fig. 5D; Fig. S2E).

### The Hippo signaling pathway is indirectly involved on the fas2^*–*^ *phenotype*

LOF conditions for Fas2 have been shown to cause over-proliferation in the follicular epithelium of the ovary and to genetically interact with *warts* (*wts*), suggesting that the Hippo signaling pathway mediates Fas2 function in this tissue [14]. We tested the interaction of *fas2* with *wts* during imaginal disc development. A reduction in half of the *wts* gene dosage rescued the wing size phenotype of the hypomorphic *fas2*^*eB112*^*/fas2*^*e76*^ combination (Fig. 5E). Since over-expression of Yorkie (*UAS-Yki*) also rescued *fas2*^*–*^ clone size (results above), the genetic interaction of *fas2* with components of the Hippo growth signaling pathway in the growing wing imaginal disc seem just opposite to that in the follicular epithelium of the ovary. To analyze the functional epistatic relationship between Fas2 and the Hippo pathway, we blocked the function of *expanded* (*ex*) and *wts* in *fas2*^*RNAi*^ FLP-OUT clones. Since both Ex and Wts are required to repress cell proliferation in imaginal discs and their LOF conditions cause overgrowth, we expected their inhibition to be epistatic to the Fas2 phenotype and cause a rescue of *fas2* clone growth (similar to Yki overexpression). However, neither condition *ex*^*RNAi*^ nor *wts*^*RNAi*^ was able to suppress the growth deficit of Fas2-deficient clones (Fig. 5C), suggesting that the Hippo pathway does not directly mediate Fas2 function in the growing imaginal discs. Since the EGFR pathway controls Yki activity [23], which in turn feeds-back on Ex expression, we studied the expression of an *ex-LacZ* reporter [24] in *fas2*^−^ cells. We detected an increase in the expression of Ex in *fas2* mutant cells (Fig. 5F), indicating an increased Yki activity. Since our previous results had shown that Fas2-deficient cells have an indirectly-induced enhanced activity in the JNK pathway (results above), and the JNK and Hippo signaling pathways interact [25], we tested if the enhanced expression of Ex in the *fas2*^*RNAi*^ condition may be a consequence of the JNK pathway increased activity. Suppression of JNK activity in Fas2-deficient cells reverted the increase in *ex-LacZ* expression (Fig. 5F). These results together with the previous ones strongly suggest that the cell competition-indirect activation of the JNK pathway in Fas2-deficient cells promotes Yki activity as a compensatory mechanism for the EGFR-dependent growth deficit.

### Fas2 physically interacts with EGFR

Our genetic data suggested that Fas2 and EGFR might physically interact at the plasma membrane. To explore this possibility we tagged the Fas2^TM^ protein isoform with a V5 epitope, and co-expressed it together with *Drosophila* EGFR (dEGFR) in HEK293 cells. Then we used an anti-V5 mouse monoclonal antibody to immunoprecipitate Fas2. A very strong anti-dEGFR immunoreactivity was pulled down from cells expressing Fas2 and dEGFR, while only background immunoreactivity was pulled down from cells that expressed dEGFR but not Fas2 (Fig. 6A). To confirm the interaction, we performed the reverse experiment. Immunoprecipitation with anti-dEGFR antibody was able to pull down Fas2 in cells that co-expressed dEGFR. However, no V5 immunoreactivity was detected when the dEGFR was not co-expressed in the cells (Fig. 6B). Together, our results demonstrate that, in the HEK293 heterologous system, Fas2 physically interacts with *Drosophila* EGFR. Since the Fas2^GPI^ isoform was sufficient to rescue the growth deficit of Fas2-deficient cells (results above), the protein interaction between Fas2 and EGFR should involve the extracellular domains of both proteins.

**Fig. 6.**
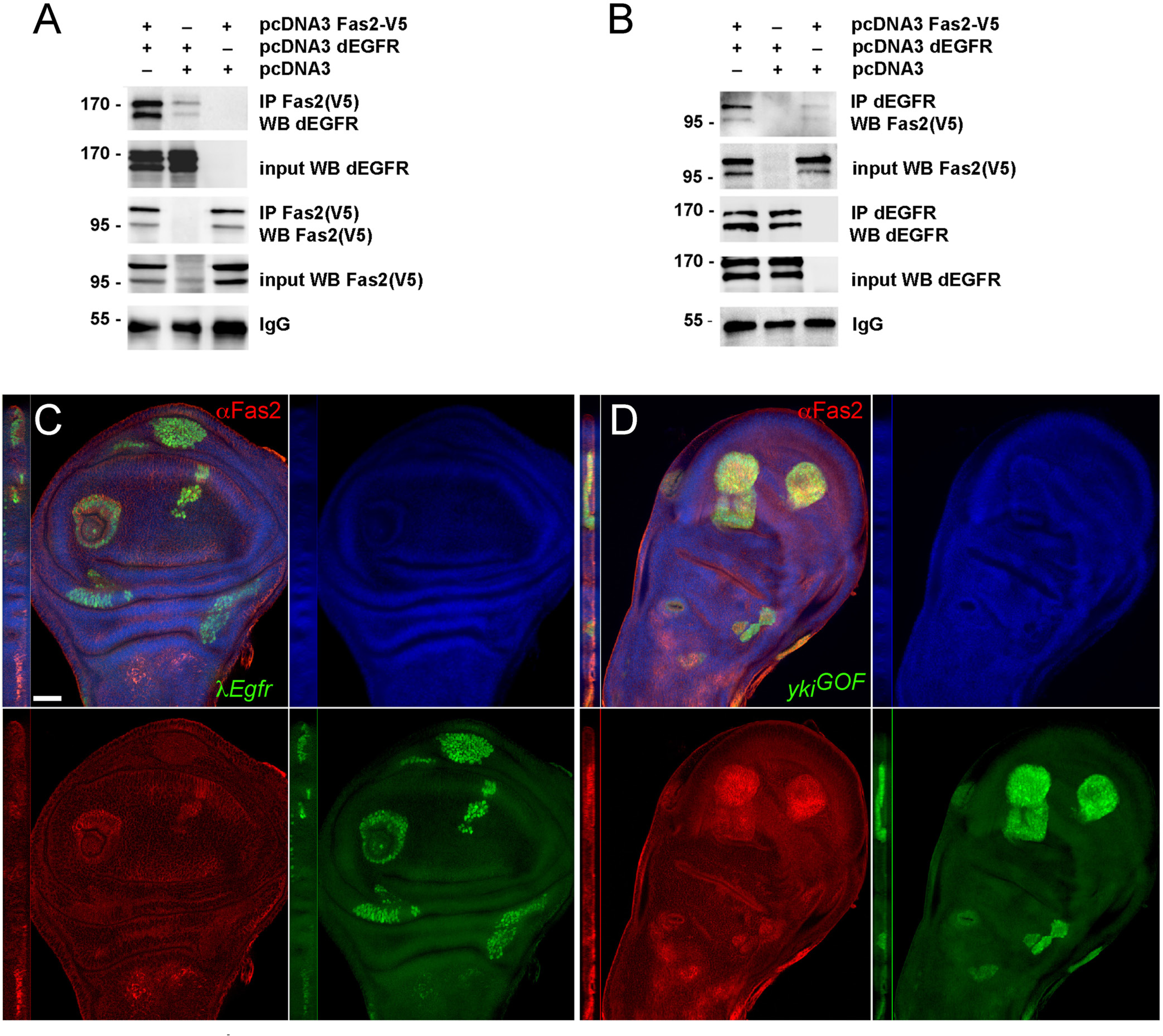
EGFR binds Fas2 and promotes its expression in imaginal discs. (A) Fas2 co-immunoprecipitates with dEGFR. Two plasmids containing the cDNA encoding for Fas2-V5 and dEGFR were transiently co-transfected into HEK293 cells (first lane). 24h later, cells were homogenized and Fas2 immunoprecipitated (IP) with the mouse anti-V5 antibody. dEGFR was strongly pulled down from these cells, while only background levels were obtained from control cells transfected with dEGFR but without Fas2 (center lane). dEGFR expression was similar in both extracts (input). The immunoblot with anti-V5 showed that the Fas2 protein was correctly expressed (input) and immunoprecipitated. IgG shows a similar load of immunoprecipitate. (B) Reverse co-immunoprecipitation. The IP with anti-dEGFR antibody pulls down Fas2 in cells co-transfected with dEGFR and Fas2, but not in cells transfected with Fas2 alone. Fas2 expression was similar in both extracts (input). Immunoblot with anti-dEGFR showed that dEGFR was correctly expressed (input) and immunoprecipitated. IgG shows a similar load of immunoprecipitate. (C) Activated-EGFR MARCM clones (*λEgfr*, labeled with GFP) show an increased expression of Fas2 (stained with the anti-Fas2 antibody MAb 1D4, red channel) in the wing imaginal disc. Similar results were obtained for the expression of activated-Ras in MARCM clones. Clones induced in 2^nd^ instar larva. Bar: 50 μm. (D) MARCM clones over-expressing Yki (*yki*^*GOF*^, labeled with GFP) display increased expression of Fas2 (in red, 1D4 antibody). Clones induced in 2^nd^ instar larva.

### EGFR and Yki activity promote Fas2 expression

EGFR LOF conditions have been shown to cause a reduction of Fas2 expression in imaginal discs [13]. We confirmed this result (Fig. S3A). In addition, we found that cell clones expressing activated-EGFR or activated-Ras (Fig. 6C) display a cell autonomous increase in the expression level of Fas2. This indicates that EGFR activity is not merely a permissive requirement for Fas2 expression in imaginal discs, but an instructive signal. Since EGFR function controls Yki activity [23], we tested if Yki over-expression could also cause a change in Fas2 expression in wing imaginal disc cell clones. We found a strong enhancement of Fas2 expression in clones over-expressing Yki (Fig. 6D). These results together with the previous ones point to the existence of a cell autonomous self-stimulating feedback loop between EGFR activity and Fas2 expression during imaginal disc growth.

**Figure S3. Fas2 dependence on EGFR function**

### Fas2 over-expression induces a non-cell autonomous restrain of epithelial growth

The previous results demonstrated that Fas2 promotes EGFR-mediated growth in the proliferative imaginal disc epithelium. To study the function in high expression conditions, we generated over-expression of Fas2 during imaginal disc development. Over-expression of Fas2 in the wing imaginal disc using the *MS1096-GAL4* driver caused a dose dependent reduction in size of the adult wing (Fig. 7A, G). Over-expression of Fas2 in *eyeless*-driven FLP-OUTs caused a reduction in size of the head capsule (Fig. 7B). Over-expression of Fas2 using *Tubulin-GAL4* or *Actin-GAL4* driver insertions caused a general reduction of body size in adult flies (Fig. S4A). Fas2 over-expression with the *engrailed-GAL4* driver caused an abnormal wing shape due to the reduction in size of the posterior compartment (Fig. S4B) without affecting cell size (Fig. S4C). This expression of Fas2^GOF^ in the posterior compartment of the wing also affected the closest region in the anterior compartment (adjacent anterior to vein 4), but not the most distant one (anterior to vein 3) (Fig. S4B).

**Fig. 7.**
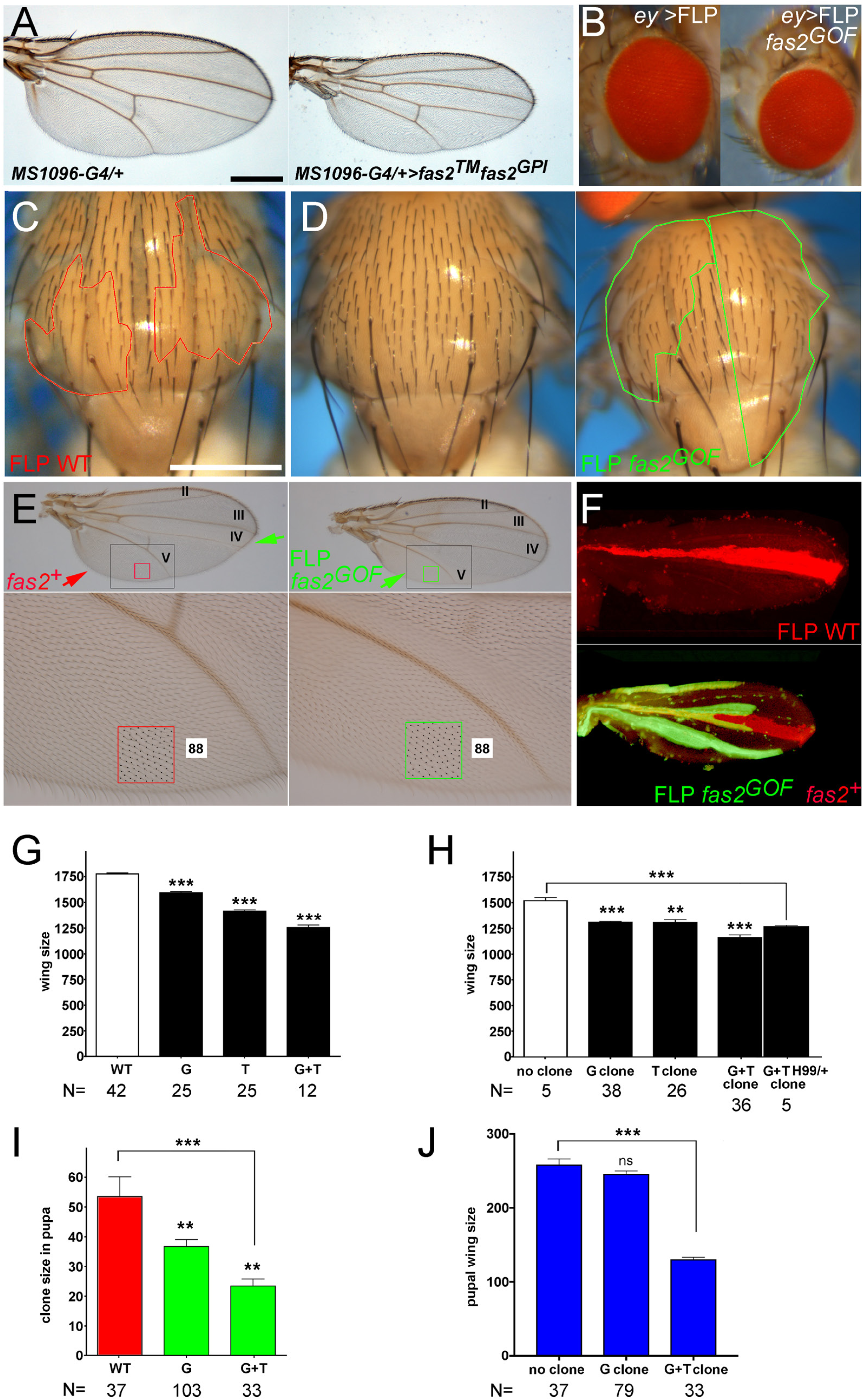
Over-expression of Fas2 causes a reduction in organ size. (A) Wild Type wing (*MS1096-GAL4/+*) in left panel. At right, over-expression of Fas2 (*fas2*^*GOF*^, *UAS-fas2*^*TM*^ *UAS-fas2*^*GPI*^*/+*) in *MS1096-GAL4/+* flies causes a general reduction of wing size. All females. Bar: 500 μm. (B) Over-expression of Fas2 (*fas2*^*GOF*^, *UAS-fas2*^*TM*^ *UAS-fas2*^*GPI*^*/+*) in *eyeless*-driven FLP-OUTs causes a general reduction of head size. (C) FLP-OUT WT notum control clones induced in 1^st^ instar larva (marked with *y* and outlined in red). All studied individuals were male. Bar: 500 μm. (D) FLP-OUT notum clones over-expressing Fas2 (*fas2*^*GOF*^, *UAS-fas2*^*TM*^ *UAS-fas2*^*GPI*^*/+*) induced in 1^st^ instar larva. Left, a control sibling without any FLP-OUT clone. Right, an individual bearing two large FLP-OUT clones (*y* cuticle, outlined in green). Note the reduction in notum size compared with either the sibling without clones or the WT control with FLP-OUT clones (in D). All studied individuals were male. (E) FLP-OUT clones over-expressing Fas2 cause a reduction of size and shape alterations in the adult wing. Top, two examples of adult male wings bearing clones that over-express Fas2^TM^ (*fas2*^*GOF*^, *UAS-fas2*^*TM*^*/+*). At the left a wing with a clone in the Anterior compartment between veins III and IV (*y* cuticle, green arrow). At right, a clone in the Posterior compartment covering the region between vein V and the posterior wing margin (*y* cuticle, green arrow). Note the local reduction in size of the regions bearing the clones compared with those same regions which do not have clones in each other wing. The panels at bottom show a magnification of the areas in the color rectangles (normal territory, *fas2*^*+*^, red arrow). The area with clone in the posterior wing at right (green frame) shows the exact same number of trichomes (each one corresponding to a single cell) than the normal region in the posterior wing at left (red frame). Note that the change in compartment size caused by the Fas2^GOF^ FLP-OUT clone is due to a reduction in the number of cells between the vein V and the wing margin. (F) FLP-OUT control (red) and Fas2^GOF^ (green) clones during pupal development. Top, a wing with a FLP-OUT WT control clone induced in 1^st^ instar larva (GFP is here shown in red). Bottom, a wing with Fas2^GOF^ clones (*fas2*^*GOF*^, *UAS-fas2*^*TM*^ *UAS-fas2*^*GPI*^*/+*, GFP in green) and an internal WT LacZ control clone (in red). The shape of the Fas2^GOF^ clones is similar to the control ones. The internal LacZ control clone runs in the dorsal and ventral surfaces of the wing (below the Fas2^GOF^ clone). Note that the pupal wing with FLP-OUT clones over-expressing Fas2 is smaller than the WT one above. (G) Quantitative analysis of *MS1096*-*GAL4* driven Fas2 GOF conditions in the adult wing. WT is *MS1096-GAL4/+* (same as in Fig. 1G), G corresponds to Fas2^GPI^ over-expression (*UAS-fas2*^*GPI*^*/+*), T to Fas2^TM^ (*UAS-fas2*^*TM*^*/+*) and G+T to Fas2^GPI^ plus Fas2^TM^ (*UAS-fas2*^*GPI*^ *UAS-fas2*^*TM*^*/+*). Wing size is wing area in μm^2^/10^3^. All individuals were female. (H) Quantitative analysis of size in adult wings bearing Fas2 GOF clones. Wings bearing FLP-OUT clones over-expressing Fas2^GPI^ (G), Fas2^TM^ (T), Fas2^GPI^ plus Fas2^TM^ (G+T) and Fas2^GPI^ plus Fas2^TM^ in a genetic background with *Df(3L)H99/+* (G+T H99) are compared with the size of wings from siblings without clones (no clone). FLP-OUT clones expressing one Fas2 GOF single insertion caused a significant reduction in adult wing size, while clones expressing two Fas2 GOF insertions displayed a much more pronounced reduction of wing size. Wing size is wing area in μm^2^/10^3^. All individuals were males. (I) Quantitative size analysis of Fas2 GOF clones in pupal wings. The size of FLP-OUT clones over-expressing Fas2^GPI^ (G) or Fas2^GPI^ plus Fas2^TM^ (G+T) (green bars) is compared with the size of WT FLP-OUT clones (WT, red bar). FLP-OUT clones with the single *UAS-fas2*^*GPI*^ insertion displayed a very significant reduction in size compared to WT FLP-OUT controls, while clones with two *UAS-fas2* insertions displayed an extremely significant reduction in size. Clone size is clone area in μm^2^/10^3^. (J) Quantitative analysis of pupal wing size in FLP-OUT Fas2 GOF clones. The size of pupal wings bearing FLP-OUT clones over-expressing Fas2^GPI^ (G) or Fas2^GPI^ plus Fas2^TM^ (G+T) is compared with the size of wings without clones from siblings (no clone). While one single *UAS-fas2* insertion (Fas2^GPI^) did not reveal a significant effect on the size of the whole pupal wing, wings bearing FLP-OUT clones expressing two *UAS-fas2* insertions displayed an extremely significant reduction in size. Wing size is wing area in μm^2^/10^3^.

**Fig. S4. Analysis of the Fas2 GOF condition in adult flies and FLP-OUT clones**

To analyze the Fas2 GOF phenotype in mosaics, we generated Fas2 GOF cell clones in the wing imaginal disc using the FLP-OUT and coupled-MARCM methods. Fas2^GOF^ clones were associated with a reduction in organ size and deformations of wing shape. Fas2^GOF^ clones in the notum that covered a significant fraction of the structure caused a striking reduction in the size of the organ compared to WT control clones or to their siblings without clones (Figs. 7C, D and S4D), suggesting that the normal territory was unable to compensate growth to attain a normal size. Adult wings with Fas2^GOF^ clones displayed strong deformations caused by a reduction in the number of cells in the compartment bearing the clone (Fig. 7E), showing that the Fas2^GOF^ effect is local. In these clones, spacing and polarity of trichomes (which mark each single epidermal cell) was normal. Wing size reduction was dependent on the level of Fas2 over-expression in the clones, and was not eliminated by suppressing apoptosis in the whole wing disc (Fig. 7H). The same dependency was obvious in pupal wings (Fig. 7F, J) compared with normal control clones (Fig. 7F). Quantification of the size of the clones in pupal wings revealed that Fas2^GOF^ FLP-OUT clones were smaller than normal in a dose dependent manner (Fig. 7I).

The analysis of Fas2^GOF^ FLP-OUT pupal wings simultaneously bearing FLP-OUT LacZ internal control clones revealed a non-cell autonomous instructive effect from the Fas2^GOF^ clones. Single control LacZ clones (without a neighbor Fas2^GOF^ clone) were significantly larger than LacZ clones growing side by side with a Fas2^GOF^ clone (Fig. S4E). Moreover, we did not detect cleaved-Caspase 3 nor *puc-LacZ* expression in these Fas2^GOF^ clones or in their neighbor cells (Fig. S4F). Thus, Fas2^GOF^ cells actively restrain the growth of their neighbors by a mechanism not involving JNK pathway signaling. The non-cell autonomous size reduction caused by the Fas2^GOF^ clones could be due to a reduction in the rate of mitosis in the clones and in the territory neighbor to the clones. Therefore, we studied the number of mitosis in wing discs bearing FLP-OUT Fas2^GOF^ and WT control clones (Fig. 8A, B). In WT controls, clones and the rim of 2-3 cells around the GFP-labeled clones displayed the same mitotic index, which was slightly but significantly higher than the general background in the disc (Fig. 8B). The mitotic index in Fas2^GOF^ clones was significantly reduced compared to the FLP-OUT WT controls. Remarkably, paired comparison of each Fas2^GOF^ clone mitotic index with that of its rim of normal cells around the clone (2-3 diameters of normal cells) (Fig. 8B) did not show any significant difference, while the background mitotic index was similar to WT controls. This indicates that the Fas2^GOF^ condition affects the normal neighbor cells around the clone. Fas2^GOF^ coupled-MARCM clones in the wing disc or pupal wing were smaller than their control twins (Fig. 8C, D). In addition, quantification of clone size in the wing discs revealed that both the Fas2^GOF^ clones and their control twins were significantly smaller than WT controls (Fig. 8E), consistent with an instructive non-cell autonomous effect of the Fas2^GOF^ clone on its neighbor cells. As with pupal wings, the size of the whole wing disc was reduced compared with WT (Fig. 8E).

**Fig. 8.**
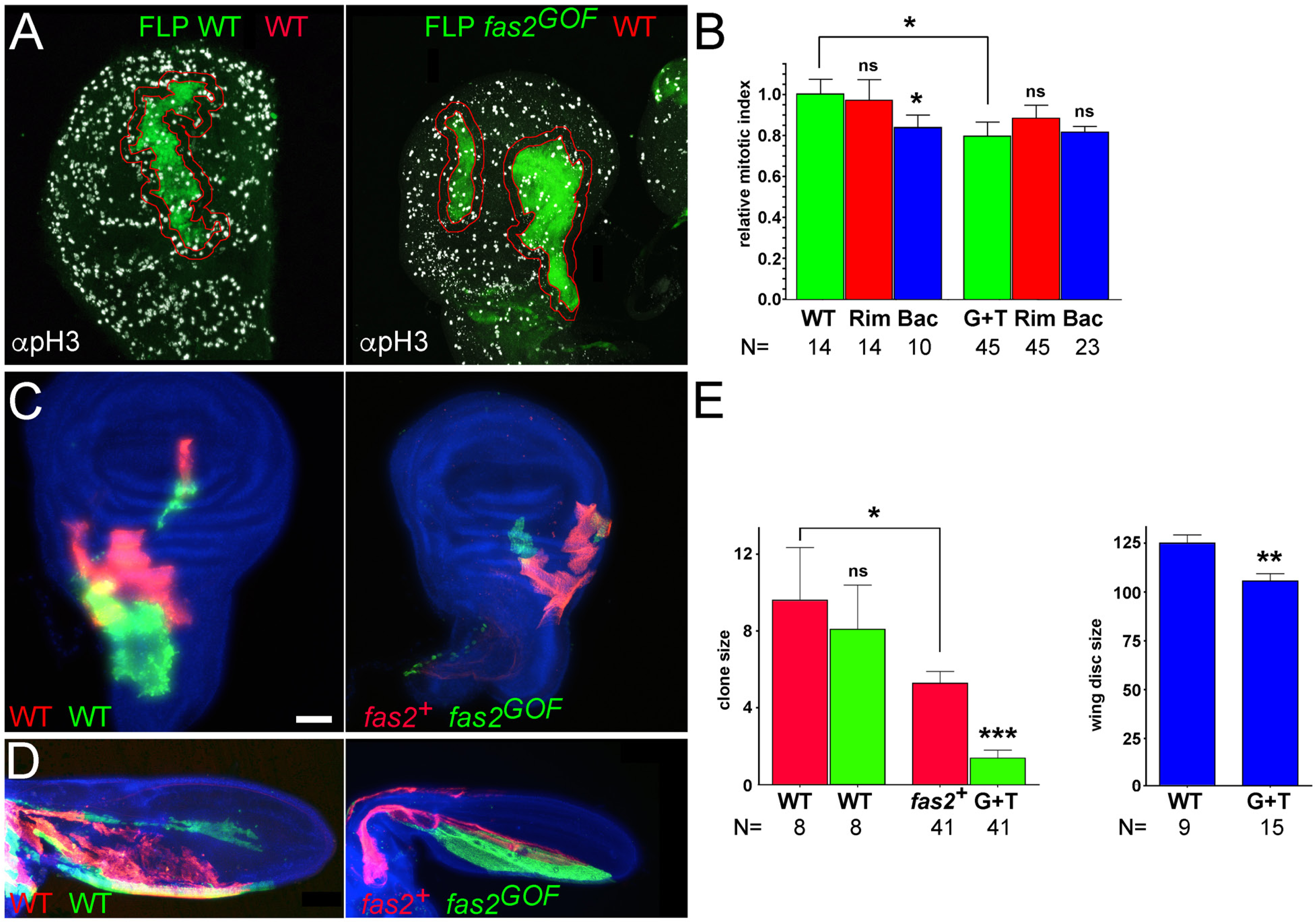
Over-expression of Fas2 causes a non-cell autonomous restraining of growth. (A) Fas2^GOF^ clones cause a non-cell autonomous reduction of the mitotic index. FLP-OUT clones expressing Fas2^GPI^ plus Fas2^TM^ (*fas2*^*GOF*^, *UAS-fas2*^*GPI*^ *UAS-fas2*^*TM*^*/+*) show a lower number of mitosis (anti-pH3 antibody, right picture) compared with control WT FLP-OUT clones (anti-pH3, left picture). We outlined a rim of 2-3 cell diameters of WT tissue around the perimeter of the clones (in red) to quantify the number of mitosis in and out of the clone (in B). (B) Mitotic index in Fas2^GOF^ FLP-OUT clones and their normal neighbor cells. FLP-OUT clones expressing Fas2^GPI^ plus Fas2^TM^ (G+T, green bar) displayed a significant reduction in the number of mitoses compared with WT controls (WT, green bar). Paired *t*-test comparison between each clone (G+T) and its neighbor cells showed that the rim of 2-3 normal cells (RIM, red bar) displayed the same mitotic index. Bac is the background relative mitotic index in the rest of the disc. Y-axis represents the number of mitosis/μm^2^ relative to WT FLP-OUT clones. (C) Coupled-MARCM analysis of the Fas2^GOF^ condition in imaginal discs. Left, coupled-MARCM WT twin clones. Right, coupled-MARCM clones in the wing disc over-expressing Fas2^GPI^ plus Fas2^TM^ (labeled with GFP) were smaller than their control twins (labeled with mtdTomato) and WT controls. Clones induced in 1^st^ instar larva. Bar: 50 μm. (D) Coupled-MARCM analysis of the Fas2^GOF^ condition in pupal wings. Left, control WT twin clones. Right, over-expression of Fas2^GPI^ plus Fas2^TM^ (*fas2*^*GOF*^, *UAS-fas2*^*GPI*^ *UAS-fas2*^*TM*^*/+*) in clones caused a reduction in pupal wing size. Clones induced in 1^st^ instar larva. (E) Left, quantitative analysis of coupled-MARCM twin clones in the wing disc. Fas2^GOF^ clones (G+T, *UAS-fas2*^*GPI*^ *UAS-fas2*^*TM*^*/+,* green bar) were smaller than their control twins (*fas2*^*+*^, red bar). In addition, these internal normal control twins were significantly smaller than WT coupled-MARCM clones (green and red bars, same as in Figure 1H), revealing the presence of a non-cell autonomous restraining of growth caused by their Fas2^GOF^ twins. Clones induced in 1^st^ instar larva. Right, quantification of wing imaginal disc size. Wing imaginal discs bearing coupled-MARCM Fas2^GOF^ clones had a significantly smaller size than WT wing disc controls. Clone size and wing disc size are expressed as area in μm^2^/10^3^.

### Fas2 functions non-cell autonomously to inhibit EGFR

To study if the non-cell autonomous functionality of the Fas2 high expression level is caused by the repression of EGFR, we first analyzed the effect of reducing the dosage of EGFR on the Fas2 GOF phenotype. A reduction in half of the *Egfr* dosage in the over-expression condition for Fas2^TM^ caused a significant enhancement of the phenotype in the wing, which became as strong as the phenotype produced by the over-expression of Fas2^GPI^ and Fas2^TM^ together (Fig. 9A). We next studied the effect of a background dosage reduction for *Egfr* in wings bearing Fas2 GOF clones. A reduction in half of the normal dose for *Egfr* caused an obvious enhancement of the phenotype without affecting the spacing or polarity of trichomes (compare Figs. 9B and 7E).

**Fig. 9.**
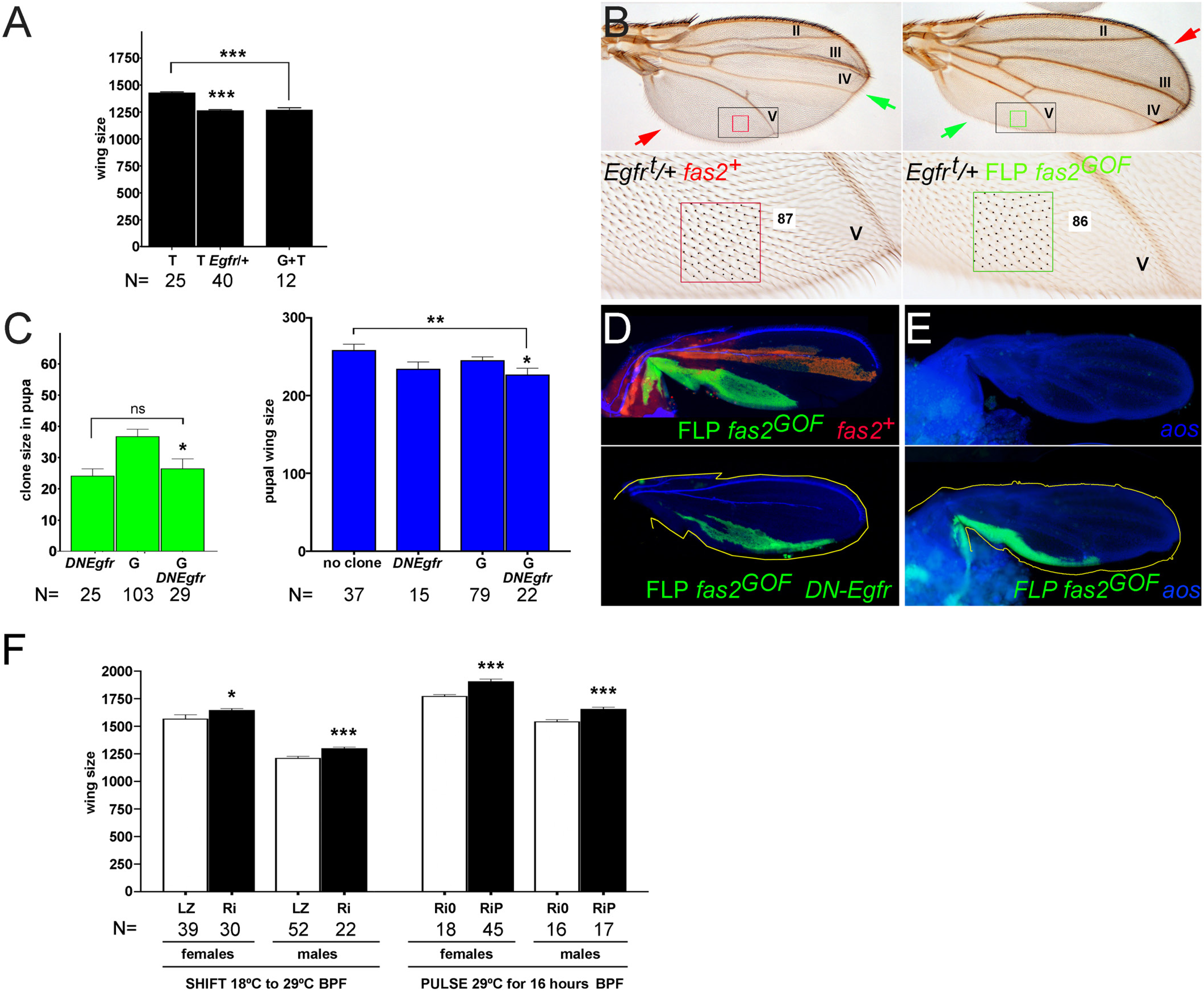
The Fas2 non-cell autonomous function inhibits EGFR and is required to limit imaginal disc growth before puparium formation. (A) A reduction in half of the dosage of *Egfr* in double heterozygous *MS1096-GAL4/+; UAS-fas2*^*TM*^*/Egfr*^*top1*^ individuals (T *Egfr/+*) causes a significant enhancement of the Fas2^TM^ GOF phenotype (T, as in Fig. 7G). The enhanced phenotype becomes similar to that obtained by the simultaneous over-expression of Fas2^GPI^ plus Fas2^TM^ (G+T, as in Fig. 7G). All individuals were female. Wing size is area in μm^2^/10^3^. (B) FLP-OUT clones over-expressing Fas2^TM^ in a genetic background with a 50% reduction of *Egfr* dosage (FLP *fas2*^*GOF*^ *Egfr/+*, green arrows). Both wings have clones in positions similar to those in Fig. 7E. The wing at the left has a clone including vein IV, and causes a reduction of size between veins II and IV, but not from vein V to the posterior margin of the wing. The wing in the right picture has a clone covering most of the Posterior compartment between veins IV and the margin, and causes the reduction in size between veins III and the posterior margin, but not between vein III and the anterior wing margin. In the *Egfr* heterozygous condition, the over-expression of Fas2 causes a stronger size reduction (compare with pictures in Fig. 7E). Moreover, size reduction is due to a lower number of cells (compare the number of trichomes between vein V and the wing margin in a normal territory, at left, with a Fas2^GOF^ territory at right). The spacing of cells (trichomes) is unchanged compared to WT territory (compare also with Figs. 1C and 7E). Clones are marked with *yellow*. Both wings are from male individuals. (C) Clone and wing size comparative analysis in pupa of FLP-OUTs over-expressing Fas2^GPI^ (G, same as in Fig. 7I, J) or Fas2^GPI^ plus a dominant negative form of EGFR, using a *UAS-DNEgfr* insertion (*DNEgfr*), (G *DNEgfr*). Left, the Fas2^GOF^ clone phenotype is not normalized, but enhanced by the expression of the hypomorphic DN-EGFR condition. The Fas2^GPI^ DN-EGFR FLP-OUT clone size is similar to DN-EGFR or to that obtained by expressing two doses of Fas2 (G+T, see Fig. 7I)). Right, the size of the wings bearing Fas2^GOF^ clones is not normalized by the expression of the hypomorphic DN-EGFR condition but further reduced, consistent with the maintenance of the Fas2^GOF^ non-cell autonomous phenotype in the presence of the cell autonomous DN-EGFR condition. Clone and pupal wing size are area in μm^2^/10^3^. (D) FLP-OUT clones labeled with GFP and over-expressing Fas2^GPI^ (FLP *fas2*^*GOF*^, top) and Fas2^GPI^ plus DN-EGFR (FLP *fas2*^*GOF*^ *DN-Egfr*, bottom) in the posterior compartment of the wing. Note that the wing bearing the Fas2^GOF^ DN-EGFR clone is smaller than the wing with the Fas2^GOF^ clone (the yellow outline corresponds to the wing above for comparison). The red clone in the top picture is a normal internal LacZ control FLP-OUT (*fas2*^*+*^). (E) The Fas2^GOF^ condition causes the non-cell autonomous restraining of growth independently of the *argos* function. The picture at top shows an *argos*^*W17E*^ mutant wing without any clones. At bottom, a FLP-OUT Fas2^GPI^ clone (FLP *fas2*^*GOF*^, labeled with GFP) in an *argos*^*W17E*^ homozygous background (the yellow outline corresponds to the wing in the top picture for comparison). (F) Quantification of wing size in flies raised at 18°C and either shifted to 29°C or exposed to a 29°C pulse for 16 hours before metamorphosis. At left (SHIFT), *Tub-GAL80*^*ts20*^*/+; hs-GAL4/UAS-fas2*^*RNAi28990*^ individuals (Ri) and *Tub-GAL80*^*ts20*^*/+; hs-GAL4*/*UAS-LacZ* controls (LZ) were raised at 18°C and transferred to 29°C on the last day of larval development. The partial mild inhibition of Fas2 expression at the end of larval life by the leaky expression of *hs-GAL4* at 29°C caused a statistically significant increase in wing size compared with the controls. At right, the partial inhibition of Fas2 expression produced by a 29°C pulse for 16 hours (PULSE) in *Tub-GAL80*^*ts10*^*/+; Tub-GAL4 UAS-GFP/UAS-fas2*^*RNAi#28890*^ individuals (Ri-16h) caused statistically significant larger wings compared with the controls (same genotype without a 29°C pulse, Ri-0h).

Fas2 LOF conditions have been reported to produce a non-cell autonomous repression of EGFR signaling during retinal differentiation [13]. Since Fas2 is maximally expressed in the differentiating retina (Fig. 1B), this function may specifically reflect its non-cell autonomous functional facet at high expression levels, as opposed to its cell autonomous role in promoting EGFR activity at lower expression levels during epithelial growth (results above). As an alternative scenario to this direct non-cell autonomous function of Fas2, we considered that Fas2 GOF conditions may create a cell autonomous over-activation of the EGFR and induce the release of a diffusible repressor of the EGFR, similar to Argos (*aos*) [26], [27], which could then indirectly mediate the non-cell autonomous effect of Fas2 GOF during imaginal disc growth. If this were the case, the non-cell autonomous repression of the EGFR in the Fas2 GOF condition would be a secondary consequence of a cell autonomous over-activation of EGFR in the Fas2 GOF clone. We tested these alternatives by expressing a weak hypomorphic dominant negative form of EGFR (DN-EGFR) in the Fas2 GOF clones. If the Fas2 GOF condition caused an over-activation of the EGFR in the Fas2 GOF cells, this increased activity should produce a partial suppression of the hypomorphic DN-EGFR cell autonomous phenotype. Reciprocally, the presence of this hypomorphic DN-EGFR component should reduce the putative increase in EGFR activity in the clone and suppress, at least partially, the Fas2 GOF phenotype. Neither was the case. Expression of the DN-EGFR was epistatic to the Fas2^GOF^ condition inside the clones, and did not reduce their effect on wing size. The results are strongly consistent with a direct EGFR repression by the Fas2 GOF (Fig. 9C, D). We also tested if Aos could be contributing in some way to the phenotype of the Fas2 GOF condition. If Aos mediated the Fas2 GOF phenotype, its deficit should be epistatic to the Fas2^GOF^ condition. This was not the case. Fas2^GOF^ FLP-OUT clones in an *aos* mutant homozygous background produced the typical reduction of wing size characteristic of Fas2^GOF^ clones (Fig. 9E). The results demonstrate that the non-cell autonomous restrain of growth produced by Fas2 GOF is not secondary to a cell autonomous increase of EGFR activity in the clone. Our combined data strongly support the idea that Fas2 behaves as a non-cell autonomous repressor of EGFR activity at high expression levels, while it functions to promote EGFR activity al low and moderate expression levels.

### Fas2 limits imaginal disc growth before metamorphosis

The previous results showed that Fas2 has two functional facets: it promotes EGFR function in a cell autonomous manner at low and moderate levels of expression (like those in proliferating imaginal discs), while it represses EGFR function in a non-cell autonomous way at high expression level (like those in the GOF chronic condition or in the differentiating retina [13]). Do these two functional facets represent different independent manners of Fas2 action in different developmental scenarios (proliferative vs. differentiating tissue)? or, do they integrate in a functional logic during imaginal disc growth? Indeed, the reciprocal dependence of Fas2 on EGFR activity during imaginal disc growth suggests that the Fas2 expression level may increase in imaginal discs during development, perhaps until reaching a threshold where the non-cell autonomous component could restrain cell proliferation. This last alternative suggested to us that a mild inhibition of Fas2 expression by the end of the larval period, still allowing enough EGFR function to fulfill its requirement for cell proliferation but moderately reducing the amount of Fas2, may cause some extra-growth.

In order to test if Fas2 restrains growth prior to metamorphosis, we used GAL80ts to specifically release the expression of *UAS-fas2*^*RNAi*^ by the end of the 3^rd^ instar larval period. We used two different strategies, either a single 16 hours 29°C pulse (restrictive temperature for GAL80ts) or a shift from 18°C to 29°C during the last 24 hours of larval development in individuals continuously raised at 18°C (permissive temperature for GAL80ts). In the first case we used *Tub-GAL80*^*ts10*^*/+; Tub-GAL4 UAS-GFP/UAS-fas2*^*RNAi#28990*^ individuals, while in the second case we used *Tub-GAL80*^*ts20*^*/+; hs-GAL4/UAS-fas2*^*RNAi#28990*^ individuals (the *hs-GAL4* insertion has a leaky expression at 29°C to drive expression of the *fas2*^*RNAi#28990*^ insertion, which is the weaker RNAi condition, see Fig. 1G). In both cases we obtained a similar result: the mild inhibition of Fas2 function prior to puparium formation caused some extra-growth compared to the corresponding controls without pulse or expressing *UAS-LacZ* (Fig. 9F). Therefore, our combined results show that the two functional facets of Fas2 operate in concert during the development of imaginal discs to promote growth during the proliferative period and to repress it by the end of larval life.

### The IgCAMs CG15630 and CG33543 mediate the non-cell autonomous function of Fas2

The previous results suggested that Fas2 control of EGFR activity switches at high expression levels. We hypothesized the possible involvement of a functional interaction that may preferentially engage at Fas2 high concentrations. Fas2 is a homophilic IgCAM that can also participate in heterophilic interactions with other IgCAMs. It has been shown that IgCAMs coded by the genes *CG15630* and *CG33543* can bind Fas2 through extracellular domain interactions [12]. The functional significance of these alternative protein interactions of Fas2 is completely unknown.

*CG15630^vbvbbbvv^ and CG33543^bgggtyu^* protein trap GFP expression was detected in all epithelial cells of imaginal discs (Fig. 10A). Interestingly, the expression within each cell does not seem to largely overlap with that of Fas2 (Fig. 10A), suggesting that these IgCAMs and Fas2 may occupy different domains at the cell membrane. To study the function of both proteins, we generated *RNAi* LOF conditions in the wing imaginal disc. Expression of *CG15630* or *CG33543 RNAi* under the control of the *MS1096-GAL4* driver caused a modest but significant reduction in wing size (Fig. 10B). Therefore, both IgCAMs seem to be required for imaginal disc growth. We then produced GOF conditions for both IgCAM genes in the wing imaginal disc. Over-expression of *UAS-CG15630* or *UAS-CG33543* insertions under the control of *MS1096-GAL4* also caused a reduction of wing size (Fig 10B). The function of CG15630 and CG33543 thus seems to be a mild parallel of Fas2 during imaginal disc growth.

**Fig. 10.**
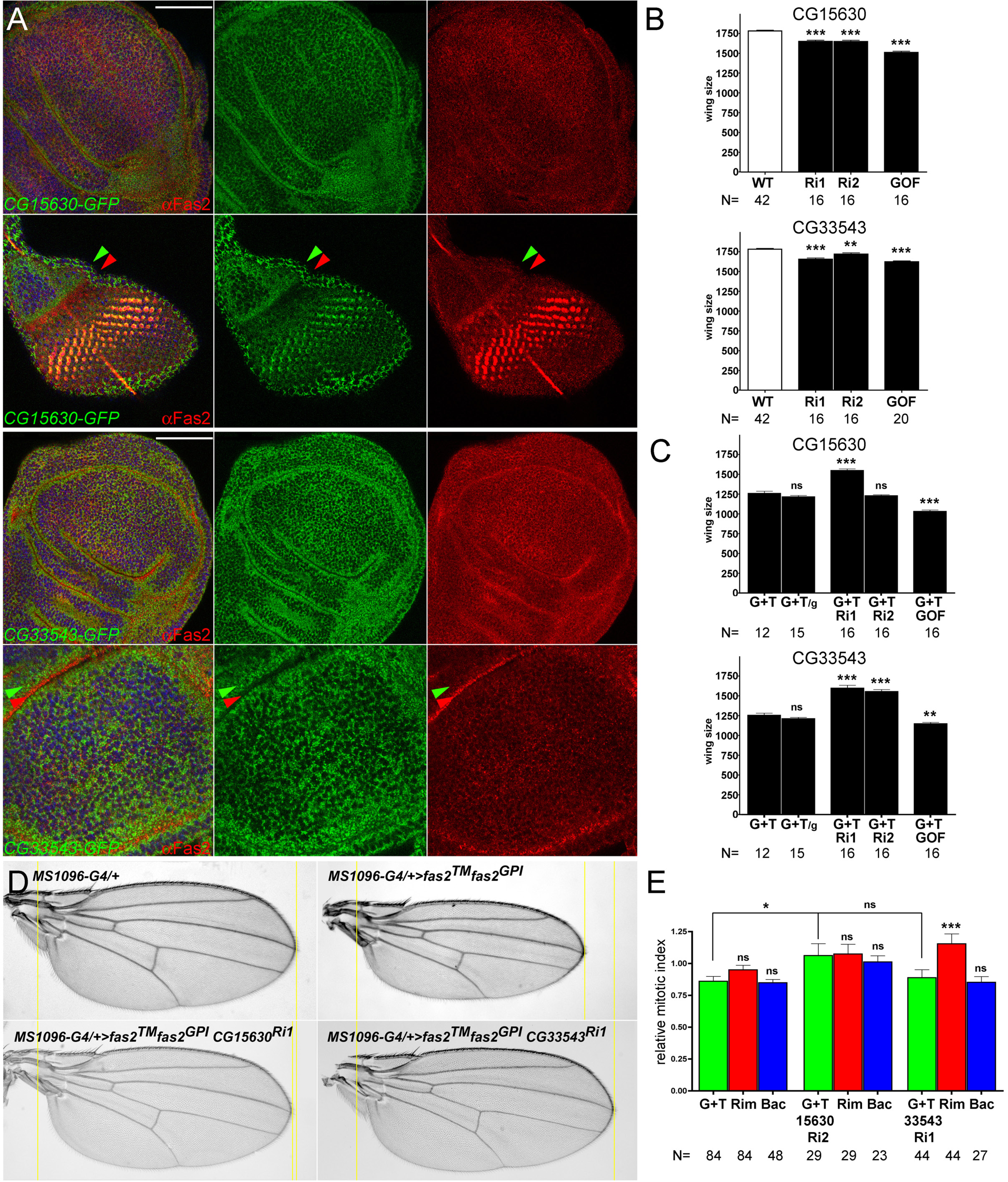
IgCAMs CG15630 and CG33543 mediate the non-cell autonomous function of Fas2. (A) Expression pattern of CG15630 and CG33543 IgCAMs in the growing imaginal discs. The protein traps CG15630-GFP-Flag #60532 (top panels) and CG33543-GFP-Flag #60531 (lower panels) are expressed in all cells of imaginal discs along with Fas2 (αFas2, 1D4 antibody). In the proliferating wing and eye imaginal discs, expression of these proteins does not exactly match Fas2 (as visualized by the little overlap between the red, Fas2, and green, protein trap, signals). In the eye antenna imaginal disc, there is obvious overlap in the non-proliferative differentiating retina (yellow cells) where Fas2 expression is maximal. The confocal section of the eye disc shows that CG15630 (green arrowhead, top panels) is expressed just adjacent to Fas2 in the same cells (red arrowhead, top panels) of the proliferative pre-Morphogenetic Furrow zone. Likewise, in the wing disc CG33543 (green arrowhead in lower panels) shows a similar expression relative to Fas2 (red arrowhead, bottom panels), being Fas2 more basally located than CG33543. Bar: 50 μm. (B) Quantitative analysis of CG15630 and CG33543 functional requirements for wing development. Above, expression of either *UAS-CG15630*^*RNAi*^ #VDRC107797kk (Ri1) or *UAS-CG15630*^*RNAi*^ #42589 (Ri2) under the control of the *MS1096-GAL4/+* driver causes a mild but highly significant reduction of wing size (WT as in Fig. 1G). Overexpression of CG15630 (in *MS1096/+; UAS-CG15630/+* females) causes a stronger reduction in wing size. Below, expression of either *UAS-CG33543*^*RNAi*^ #VDRC32576 (Ri1) or *UAS-CG33543*^*RNAi*^ #VDRC40821 (Ri2) under the control of the *MS1096-GAL4/+* driver causes a mild but significant reduction of wing size (WT as in Fig. 1G). Overexpression of CG33543 (in *MS1096/+; UAS-CG33543/+* females) also causes a reduction in wing size. (C) Quantitative analysis of the Fas2^GOF^ genetic interaction with CG15630 and CG33543. The Fas2^GOF^ (G+T, *MS1096-GAL4/+; UAS-fas*^*GPI*^ *UAS-fas2*^*TM*^*/+*) wing phenotype is not modified by the introduction of and additional *UAS* (*UAS-GFP*) control insertion (G+T/g). Above, expression of *CG15630*^*RNAi*^ #VDRC107797kk (G+T Ri1) in *MS1096-GAL4/+; UAS-fas2*^*GPI*^ *UAS-fas2*^*TM*^*/UAS-CG15630*^*RNAi*^ females causes a strong suppression of the Fas2^GOF^ wing phenotype. Over-expression of *CG15630* in *MS1096/+; UAS-fas2*^*GPI*^ *UAS-fas2*^*TM*^*/ UAS-CG15630* females (G+T GOF) causes a reciprocal enhancement of wing size reduction. Below, expression of *CG33543*^*RNAi*^ #VDRC32576 (G+T Ri1) or *CG33543*^*RNAi*^ #VDRC40821 (G+T Ri2) in *MS1096-GAL4/+; UAS-fas2*^*GPI*^ *UAS-fas2*^*TM*^*/UAS-CG33543*^*RNAi*^ females rescues the Fas2^GOF^ wing phenotype. Over-expression of *CG33543* in *MS1096/+; UAS-fas2*^*GPI*^ *UAS-fas2*^*TM*^*/ UAS-CG33543* females (G+T GOF) causes a reciprocal enhancement of the Fas2^GOF^ non-cell autonomous phenotype. (D) Epistatic relationship of Fas2^GOF^ with CG15630 and CG33543. Inhibition of *CG15630* or *CG33543* expression blocks the production of the Fas2^GOF^ non-cell autonomous phenotype in *MS1096-GAL4/+; UAS-fas2*^*GPI*^ *UAS-fas2*^*TM*^*/UAS-CG15630*^*RNAi*^ or *MS1096-GAL4/+; UAS-fas2*^*GPI*^ *UAS-fas2*^*TM*^*/UAS-CG33543*^*RNAi*^ females. (E) Quantitative analysis of mitotic index in the genetic interaction of Fas2^GOF^ with CG15630 and CG33543. Over-expression of Fas2 (*UAS-fas2*^*GPI*^ plus *UAS-fas2*^*TM*^, G+T) in FLP-OUT cell clones causes a reduction of the mitotic index in the clone and in the rim of 2-3 normal neighbor cells around the clone. Co-expression in these FLP-OUT clones of *CG15630*^*RNAi*^ #42589 (G+T 15630Ri2) increases the mitotic index in both the clone and the rim of normal cells around the clone. Co-expression in the Fas2^GOF^ clones of CG33543RNAi #VDRC32576 (G+T 33543Ri1) specifically rescues the non-cell autonomous effect in the rim of normal cells around the clone. Y axis: mitotic index relative to WT clones.

To study the possible participation of these IgCAMs in mediating the non-cell autonomous function of Fas2, we expressed their *RNAi* insertions in the wing imaginal disc of Fas2^GOF^ individuals. Remarkably, expression of different *CG15630*^*RNAi*^ or *CG33543*^*RNAi*^ lines completely suppressed the Fas2^GOF^ phenotype in adults (Fig. 10C, D). We then combined Fas2^GOF^ with the CG15630 or the CG33543 GOF condition, and we obtained an enhancement of the Fas2^GOF^ non-cell autonomous phenotype in both cases (Fig. 10C). To test if the suppression of the non-cell autonomous Fas2 phenotype affected the mitotic index in imaginal discs, we generated FLP-OUT clones over-expressing Fas2 along with *CG15630*^*RNAi*^ or *CG33543*^*RNAi*^. A general increase in the mitotic index, autonomous and non-cell autonomous, occurred for the *Fas2*^*GOF*^ *CG15630*^*RNAi*^ combination, while a specific non-cell autonomous correction of the mitotic index in the rim of 2-3 normal neighbor cells around the clone occurred for the *Fas2*^*GOF*^ *CG33543*^*RNAi*^ clones (Fig. 10E). The results show that CG15630 and CG33543 IgCAMs are required in the same cell than Fas2 to mediate its non-cell autonomous function (Fig. 11A). In summary, Fas2, CG15630 and CG33543 IgCAMs function in parallel to promote imaginal disc growth. In addition, Fas2 engages in interactions with CG15630 and CG33543 at high expression levels to produce a non-cell autonomous restrain of growth.

## Discussion

We have characterized the function of the *Drosophila* IgCAM protein Fas2 in the imaginal disc growing epithelium. Fas2 is the *Drosophila* homolog of vertebrate NCAM, and, as its vertebrate counterpart, acts as a homophilic CAM during axon and synapse growth. Interestingly, a general observation is that reduced or enhanced levels of CAM-mediated contact reduce neuronal process growth, following the general principle that just a correct amount of adhesion is required for optimal neuronal-process formation (see [28]). Fas2 promotes maximal synaptic growth at an expression level of 50%, with lower or higher levels causing a smaller number of synaptic boutons [29]. Expression of Fas2 at the synapse has been shown to depend on the activity of Ras [30], a main EGFR effector. In addition to its established role as recognition/adhesion mediator, Fas2 has been shown to promote EGFR function during axon extension [8]. Conversely, Fas2 has been previously reported to function as an EGFR repressor during retinal differentiation [13], as vertebrate NCAM can do during axon extension [31].

Our analyses during imaginal disc growth reveal that the role of Fas2 in repressing EGFR function occurs at the highest expression levels of the protein, like those attained during the stop of cell proliferation at the Morphogenetic Furrow and the initiation of retinal differentiation. In contrast, Fas2 promotes cell proliferation and epithelial growth in imaginal discs at low and moderate expression levels. Accordingly, Fas2 is expressed in all cells of the proliferating epithelium of imaginal discs at a lower level than in the differentiating imaginal structures (retina, sensory organ precursor cells and veins in the wing disc). Our LOF analysis shows that imaginal organ growth is titrated by the level of Fas2 function. Fas2 LOF genetic mosaics reveal that imaginal disc cell clones deficient for Fas2 grow slower than their normal twins, showing a cell autonomous requirement of Fas2 for cell proliferation. This cellular requirement is dependent on the extra-cytoplasmic part of Fas2, since it can be corrected simply by expressing the Fas2^GPI^ isoform in the Fas2-deficient cells. This requirement reflects a dependence of proliferation on the Fas2 homophilic interaction between cells (rather than a requirement for the simple presence of the protein), since both Fas2-deficient cell clones and their closest Fas2-expressing neighbor cells have a reduced mitotic index. In contrast to the neighbor cells lacking the homophilic interaction with the *fas2*^*–*^ clone, those normal cells not directly confronting *fas2*^*–*^ clones are able to compensate for the loss of growth to attain a correct organ size. The loss of growth in Fas2-deficient clones correlates with an over-activation of the JNK signaling pathway, as revealed by the strong expression of the reporter *puc-LacZ* in cells or compartments deficient for Fas2. However, this JNK over-activation does not cause the phenotype of Fas2-deficient clones, since blocking cell death or JNK activity does not correct the loss of growth in *fas2*^*–*^ clones. The increased activity of the JNK signaling pathway in the *fas2*^*–*^ cells depends on the growth rate of the Fas2-normal background, as revealed by its reversion when the Fas2-deficient cells are growing in a normal but slow proliferating cellular background. Our data show that JNK activation in Fas2-deficient cells indirectly results from a cell interaction mechanism during cell competition that tends to compensate growth rate between cells. In addition, we found that normal Puc gene dosage is critical for Fas2-deficient cell survival. Compartments deficient for Fas2 can proliferate (slower than normal) and differentiate in adults, but a reduction in 50% the normal dosage of Puc causes widespread cell death and the collapse of the compartment. This indicates that the regulatory feedback between JNK and Puc plays a pivotal role in this compensatory mechanism during cell competition.

The requirement for Fas2 during imaginal disc growth reflects a function to promote EGFR activity. Fas2-deficient imaginal discs show a reduced level of MAPK activation, and *fas2*^*–*^ clones show a reduced mitotic index. In addition, LOF and acute GOF (but not chronic GOF) conditions of *fas2* display phenotypes reminiscent of *Egfr* LOF and GOF conditions respectively. The genetic interaction epistasis of *fas2* and *Egfr* LOF mutant combinations shows that Fas2 and EGFR function in the same developmental pathway during imaginal disc growth. We have also found that Fas2 physically binds EGFR. Moreover, the cell autonomous slow growth of *fas2*^*–*^ clones is rescued by the expression of activated-EGFR or activated forms of its downstream effectors: Raf, Ras or PI3K. It should be noted that although both the EGFR and the FGFR mediate Fas2 function in growing sensory axons [8], the expression of activated-FGFR, which shares most of the downstream effectors with the EGFR [32], is unable to produce any rescue in growing imaginal discs. This reveals a high degree of specificity in the molecular interactions that mediate Fas2 function in different tissues during development. It is very significant that EGFR activity in turn drives the cell autonomous expression of Fas2 in imaginal discs. We have shown that EGFR LOF and GOF conditions cause corresponding changes in the expression level of Fas2, therefore this is an instructive function of EGFR. Thus, the combined results of the Fas2 LOF analysis reveal the existence of a cell autonomous, self-stimulating, positive feedback loop between Fas2 and the EGFR signaling pathway during imaginal disc growth (Fig. 11A).

We also analyzed the contribution of the Hippo signaling pathway to the Fas2 function during imaginal disc cell proliferation. The Hippo pathway is critical to control cell proliferation during imaginal disc development [33] and mediates JNK-dependent cell competition interactions [34] [35]. Moreover, *fas2* LOF mutations have been shown to genetically interact with *wts* in the follicular epithelium of the ovary [14]. Fas2 function in this epithelium was proposed to involve an activation of the Hippo pathway (a repression of growth), since *fas2* LOF mutants produced a phenotype of extra cell proliferation consistent with an insufficiency in this pathway. In contrast, we have detected a genetic interaction between *fas2* and *wts* in the imaginal disc epithelium just opposite to that in the follicular epithelium. Our results indicate that this action of Fas2 on the Hippo pathway is indirect in the growing imaginal discs. First, we do not detect any rescue of Fas2-deficient cell proliferation when blocking the Hippo pathway by the expression of *ex*^*RNAi*^ or *wts*^*RNAi*^, contrary to what would be expected if Ex and Wts directly mediated Fas2 function during imaginal disc growth. Second, although we indeed detected an increase in expression of the reporter *ex-LacZ* in Fas2-deficient cells (indicating an enhanced Yki expression), this is reverted to normal when we simultaneously inhibit the indirect activation the JNK pathway in these cells. Third, our results do show that over-expression of the Hippo pathway effector Yki can rescue the Fas2-deficient clone growth, but Yki has also been shown to be regulated by EGFR and JNK activity [23] [25]. Therefore, Yki involvement in mediating the *fas2*^*–*^ phenotype most probably reflects its dependence on both, a direct effect of Fas2 deficit on EGFR activity and an indirect compensatory effect via JNK action on the Hippo pathway.

In addition to its cell autonomous function to promote EGFR-dependent cell proliferation in growing imaginal discs, Fas2 harbors a complementary non-cell autonomous functional facet to repress growth at high expression levels. We have found that Fas2 GOF conditions cause a dose dependent reduction of wing size. The phenotype of these Fas2 GOF conditions is highly sensitive to the dose of the *Egfr* gene. Interestingly, *fas2* LOF mutations have been shown to cause over-proliferation in the follicular epithelium of the ovary [14], and to produce a non-cell autonomous derepression of the EGFR in differentiating cells of the retina [13]. In both situations Fas2 is expressed at high levels. Now, our analysis of Fas2 GOF conditions in growing imaginal discs shows that Fas2 is not a mere permissive factor for cell proliferation. High expression levels of Fas2 actively repress imaginal disc cell proliferation in a non-cell autonomous manner through the EGFR. This repression is local, since preferentially affects the neighbor normal cells of Fas2^GOF^ clones. We do not know how far this non-cell autonomous function of Fas2 extends, but reaches beyond the first row of cells contacting the Fas2-deficient clone. The result that wings with large Fas2^GOF^ clones display strong alterations of shape reveals that the effect is most prominent for those normal cells closer to the clone. Inhibition of Fas2 expression in the posterior compartment of the wing causes a reduced growth in the posterior compartment and in the region between veins 3 and 4 in the anterior compartment, but does not affect growth from vein 3 to the anterior margin of the wing. A long-range effect of Fas2^GOF^ may result from the involvement of cytonemes [36] in Fas2 cell to cell communication in imaginal discs. Alternatively, or at the same time, it may result from the shedding of the Fas2 extra-cytoplasmic domain, as it has been shown for NCAM [37]. In either case, note that the Fas2^GOF^ active repression on the neighbor cells reduces the size of the twin normal clones, thus it is notably different from that passive effect resulting from the loss of homophilic interaction between contacting cells in Fas2-deficient clones (which just affects the normal cells contacting the clones but not the size of the control twins). Importantly, the non-cell autonomous repression of EGFR function is not a secondary effect of a high EGFR activity in the Fas2 GOF cells and the corresponding secretion of some putative feedback repressor, since a simple reduction in half of EGFR or the expression of a mild DN-EGFR condition in the clones enhances the phenotype instead of suppressing it. Rather, this is strongly consistent with Fas2 bearing an alternative functional facet at high expression level to actively repress EGFR in neighbor cells (Fig. 11A).

**Fig. 11.**
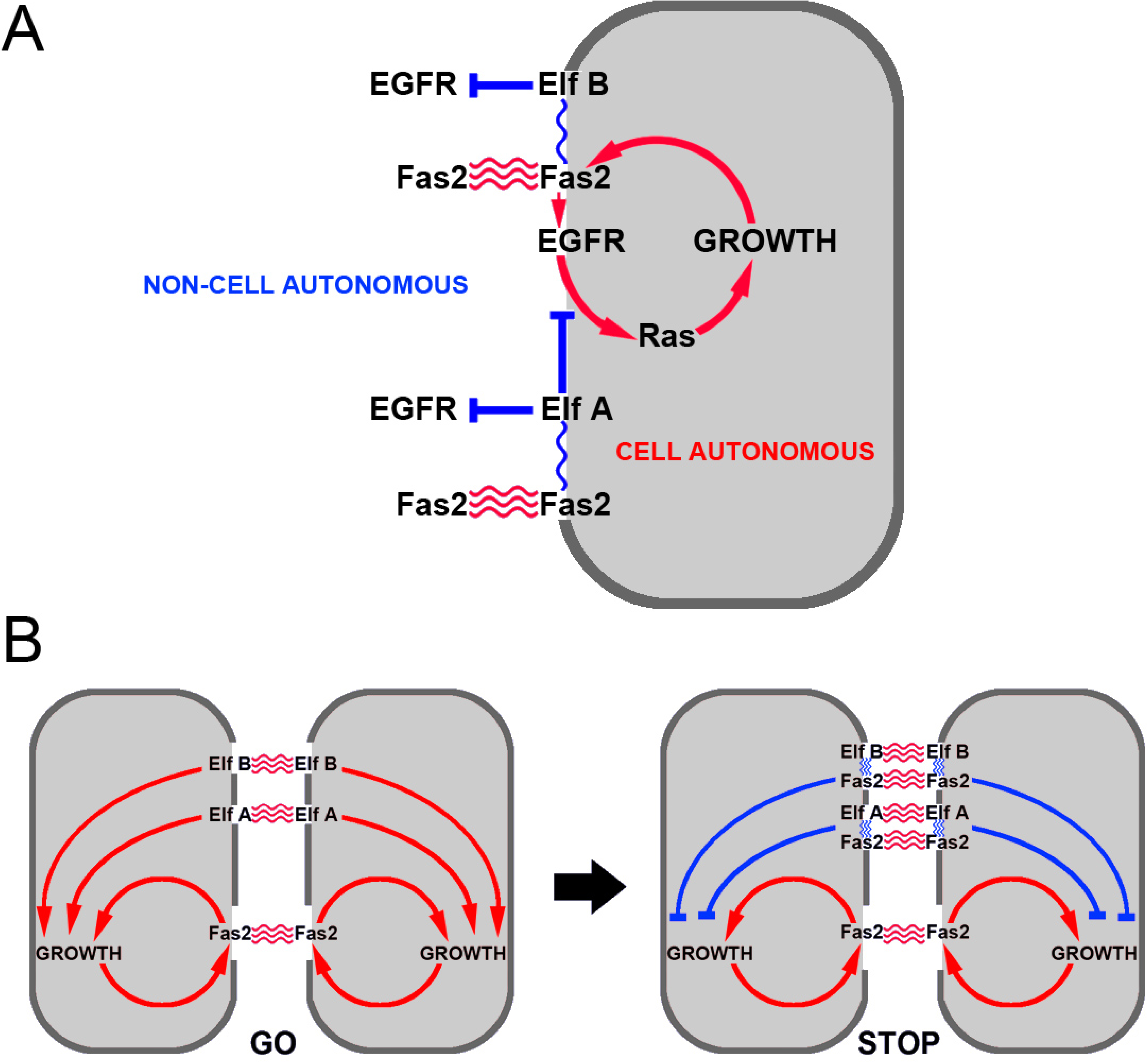
Fas2 functions as an expression level switch of EGFR during imaginal disc growth. (A) Summary of functional genetic interactions found in this study. A cell autonomous positive feedback between the homophilic-activated Fas2 and the EGFR function is shown in red. Fas2 homophilic adhesion (red) is autonomously required to promote EGFR function (through direct physical interaction). EGFR activity in turn promotes Fas2 expression creating a cell autonomous positive feedback loop. A non-cell autonomous repression of EGFR mediated by the heterophilic interaction with ElfA (Expression level fencer A, CG15630) and ElfB (Expression level fencer B CG33543) is shown in blue. (B) Hypothetical model for a Fas2-level switch mechanism controlling cell proliferation and organ size in imaginal discs. Growth control in proliferating epithelia of imaginal discs would depend on cell autonomous (in red) and non-cell autonomous (in blue) functions of Fas2, ElfA and ElfB. Fas2, ElfA and ElfB would engage in homophilic interactions that promote cell proliferation. In addition, the three IgCAMs would participate in alternative heterophilic interactions to repress EGFR in neighbor cells. A differential binding affinity of the homophilic vs. the heterophilic interactions and their differential expression between cells could modulate the rate of cell proliferation in a way similar to the proposal of the Entelechia model [6]. The absolute amount of Fas2 could be instructive to switch off growth once reaching a certain threshold level, acting as an expression level functional switch.

The dissection of the cell autonomous and non-cell autonomous functional facets of Fas2 suggests that this protein is both a cell autonomous EGFR activator at low and moderate expression levels, and possibly a heterophilic ligand for another cell membrane protein that helps Fas2 to switch to the non-cell autonomous function at high expression levels. This heterophilic interaction could mediate the repression of EGFR or its signaling pathway in neighbor cells. The two functional facets of Fas2 operate in a concerted manner during imaginal disc growth. We have shown that inhibition of Fas2 expression during larval development impedes EGFR-dependent growth, but a moderate decrease of expression just prior to puparium formation causes some extra-growth. Thus, by the end of the proliferation period in imaginal discs, the Fas2 integrated functional output has switched from promoting EGFR-dependent growth to repressing it.

CG15630 and CG33543 IgCAMs have been shown to heterophilically bind Fas2 through extracellular domain interactions [12]. Therefore, these proteins are good candidates to interact with Fas2 in high expression situations. Analysis of the expression pattern of CG15630 and CG33543 shows that during imaginal disc growth these proteins are expressed in the same cells that Fas2, but their domains do not exactly match with Fas2 at the cell membrane. The function of both proteins seems to parallel that of Fas2 during imaginal disc development: they are required for epithelial growth and their over-expression is able to repress it (although they seem less important than Fas2 to sustain cell proliferation). Our results show that both proteins mediate the Fas2 non-cell autonomous function. Their LOF condition suppresses Fas2^GOF^ while their GOF condition enhances it. Remarkably, both proteins are required in the same cell than Fas2 to produce the non-cell autonomous function, although they seem to have a different way of action. While CG33543 seems to act exclusively in a non-cell autonomous manner, CG15630 seems to act both cell autonomously and non-cell autonomously (Fig. 11A). Given the role of these proteins to limit the action of Fas2 on EGFR function, we propose to name them as Fas2 Expression level fencer A (Elf A, CG15630) and Expression level fencer B (Elf B, CG33543). The partially matching pattern of expression of these proteins and Fas2 in the same cells is strongly consistent with the idea that the heterophilic interactions between these proteins could start to occur (or become more frequent) in the high Fas2 expression situation. In the simplest hypothetical scenario, heterophilic interactions between different homophilic IgCAMs could simultaneously switch the function to mutual inhibition in all of them to terminate growth in imaginal discs (Fig. 11B).

We hypothesize that the integration of the Fas2, ElfA and ElfB cell autonomous and non-cell autonomous functional facets may provide a simple mechanism to drive and compute cell proliferation in the growing epithelium (Fig. 11B). The relative amount of these IgCAMs between cells would be critical to titrate the homo-vs. heterophilic interaction balance towards promoting growth or restraining it. Amounts of Fas2, ElfA and ElfB lower than a given threshold would favor homophilic interactions and promote EGFR/Ras pathway activity, producing in turn an increase of Fas2 expression due to its positive feedback with EGFR. This loop would be self-stimulatory until reaching over that Fas2-dependent threshold, when the heterophilic interactions would start to restrain EGFR activity and cell proliferation in a non-cell autonomous manner. In this Fas2 gain-dependent growth model, Fas2 acts as a sensor in a growth level switch, and the final size of the organ is an emergent property that relies on the difference between the affinities of the homo-and heterophilic interactions of Fas2, ElfA and ElfB. Future work should reveal if ElfA and ElfB function as homophilic IgCAMs in a way similar to Fas2. The proposed mechanism places cell-to-cell communication codes (the affinities of growth promoting vs. growth restraining IgCAM interactions) as the driving force behind the control of growth and shape. In our view growth and shape regulation is fundamentally a negotiation between cells.

During developmental growth and regeneration of epithelial organs, cell recognition is thought to drive intercalary cell proliferation. Cell recognition is also important to control axon growth and synapse formation and plasticity. Fas2 and other CAMs that could engage in alternative homo-vs. heterophilic interactions are good candidates to constitute expression level switches for the control of size. These molecules couple highly specific cell recognition with the activation and repression of signaling mechanisms that control growth. Thus, CAM expression-level switches may constitute universal mechanisms to concert growth and size control during epithelial organ development, axon extension and synapse formation.

## Materials and Methods

### Drosophila strains and genetic crosses

A description of the different *Drosophila* genes and mutations can be found at FlyBase, *www.flybase.org*. *Drosophila* stocks were obtained from the Bloomington Stock Center and Vienna Drosophila Research Center. Plasmids containing full length cDNAs of *CG15630* and *CG33543* were purchased from the Drosophila Genomics Resource Center, and the corresponding *UAS-CG15630 and UAS-CG33543* constructs and transformant lines were obtained from Best Gene, Inc., California. The different strains used in the mosaic analyses can be found in the Supplementary Materials and Methods. Clones were induced by 1-hour heat-shock at 37° C, for MARCM and coupled-MARCM analyses, and a 10 minutes heat-shock, for FLP-OUT clones, during 1^st^ (24-48 hours after egg laying, AEL) or 2^nd^ (48-72 hours AEL) larval instars. All genetic crosses were maintained at 25±1°C in non-crowded conditions. To obtain *fas2*^*–*^ *Minute*^*+*^ *puc-LacZ* clones, we crossed *fas2*^*eB112*^ *FRT19A; puc-LacZ/hs-fas2*^*TM*^ males with *Ubi-GFP M(1)O*^*sp*^ *FRT19A/FM7; hs-FLP* females and followed a typical heat-shock protocol to induce clones in 1^st^ and 2^nd^ instar larva. *hs-fas2*^*TM*^ and *puc-LacZ* dissected imaginal discs were identified by the normal expression of *puc-LacZ* in the peripodial membrane at the base of the wing disc.

### Fas2 and EGFR co-immunoprecipitation

The pcDNA3 construct encoding for the *Drosophila* EGFR (*DER 2*) was a gift of Prof. E. Schejter (Dept. of Molecular Genetics, The Weizmann Institute of Science, Rehovot, Israel). The construct *pOT2-Fas2* was obtained from the *Drosophila* Genomics Resource Center. The cDNA encoding for *Drosophila fas2* was subcloned into the *pcDNA3.2/V5/TOPO* vector (Invitrogen). Human embryonic kidney cells (HEK293) were cultured in Dubecco′s modified Eagle′s medium (DMEM) containing 10% of fetal bovine serum. Cells were plated at sub-confluence, and twenty hours later transfected with the indicated plasmids using lipofectamine 2000 following the manufacturer recommendations. To improve Fas2 expression, cells were incubated for 12h with 2 μM MG132. 24h post-transfection cells were lysed (lysis buffer: 50mM Tris, 150mM NaCl, 2mM EDTA, protease inhibitor cocktail-Complete Mini, Roche-, 0.5% Triton X-100) and incubated 1h on ice. Cell lysate was centrifuged 10min at 4°C, and an aliquot of the supernatant was kept aside on ice (“input”). Protein A-Sepharose beads (GE Healthcare) were loaded with rabbit anti-V5 tag ChIP grade (Abcam; ab9116) or mouse anti-*Drosophila* EGFR (C-273) antibody (Abcam; ab49966) for 1h at room temperature and washed 3 times with PBS. Cell lysate supernatant was mixed with antibody-loaded beads, and incubated 3h on ice, with mild shaking. Beads were washed 4 times with ice-cold PBS, resuspended in SDS sample buffer, boiled 5min, and submitted to SDS-PAGE in a 7% acrylamide gel. The proteins were transferred to nitrocellulose membrane (Protran, Whatman GmbH). The membrane was blocked with 5% BSA in TBS containing 0.1% Tween and incubated with the indicated antibody in blocking buffer overnight at 4 °C. The membrane was then washed three times with TBS containing 0.1% Tween, and the corresponding secondary antibody (horseradish peroxidase-conjugated: SIGMA anti rabbit IgG-HRP-A9169-and anti-mouse IgG-HRP-A4416-) was applied at 1:2000 dilution in TBS containing 0.1% Tween for 2h at room temperature. Immunoreactivity was detected using ECL Plus detection reagent (GE Healthcare).

### Immunohistochemistry and data acquisition

We used the following primary antibodies: anti-Futsch MAb 22C10 for PNS neurons, anti-Fas2 MAb 1D4 and anti-*β*gal MAb 40-1 from the DSHB (University of Iowa); anti-*β*gal rabbit serum from Capel; rabbit anti-cleaved Caspase3 and anti-phospho-Histone H3B from Cell Signaling Technology, and anti-activated MAPK from Sigma. Staining protocols were standard according to antibody specifications. Images of adult nota and wings were acquired at 40X, pupal wing images at 100X, and imaginal discs at 200X. These images were used for surface measurements in pixels (pxs) and expressed as μm^2^ according with the calibration of the microscope objective and digital camera used. Measured wing area corresponded to the region from the alula to the tip of the wing. Confocal images were obtained in a Leica DM-SL upright microscope, or in a Nikon Eclipse i80 microscope equipped with an Optigrid Structured Light System. Images were processed using Volocity 4.1-6.1 software, Improvision Ltd, Perkin-Elmer.

### Statistical analysis

To calculate the mitotic index in the clone, rim and rest of the imaginal disc, we outlined each GFP clone with the selection tool in Photoshop. To obtain the rim area of normal tissue corresponding to some 2-3 cell diameters around the GFP clone, we extended the selection area of the GFP signal by 25 pixels out of the clone border and removed the area of the clone itself. The area between this rim and the border of the disc defined the rest of the tissue. We counted the number of pH3 positive spots in each area, divided this number by the area in μm^2^ according with magnification and camera resolution. All values were then expressed as ratio to the mitosis/area value of the WT control clones. We used paired Student’s *t*-test (two-tailed) when possible, as it is the most powerful and stringent test to compare differences between clone twins and between each clone, its rim and its imaginal disc background. In other cases, to compare values between different populations, we used unpaired Student’s *t*-test (two-tailed) when variances were similar and fitted a Gaussian distribution, Student’s *t*-test with Welch’s correction when the variances were different and the Mann-Whitney test when the distribution of values was not gaussian. Error bars in Figures are SEM. Significance value: * P<0.05, ** P<0.01 and *** P<0.001. Statistical software was Prism 4.0c and 7.0d, GraphPad Software, San Diego California USA, *www.graphpad.com*.

## Author contribution

Genetics: EV and LGA; coIPs: ED, CGM, JGS, FG; Conceptual design: LGA; Experimental design: LGA (genetics) and HC (coIPs); Manuscript preparation and writing: LGA

## Acknowledgements

We thank S. Campuzano and P. Martin for the *w M(1)O*^*sp*^ *FRT18A/FM7; hsp70-flp; Dp(1;3)A59*/*TM6B* and the *puc-LacZ* strains, and E. Schejter for the *pcDNA3-EGFR* plasmid. We are grateful to S. Baars for technical help in some experiments and D. Ferres-Marco for thoughtful comments on the manuscript. Most primary antibodies were obtained from the Developmental Studies Hybridoma Bank.

## Financial Disclosure

This work was supported by SAF2004-06593 from MCYT, PROMETEO 2008-134, PROMETEO II 2013-001 from Generalitat Valenciana, BFU2016-76295-R from MINECO and a MCyT FP-2001-2181 predoctoral fellowship to EV. The funders had no role in study design, data collection and analysis, decision to publish, or preparation of the manuscript.

